# Tracking single-cell gene regulation in dynamically controlled environments using an integrated microfluidic and computational setup

**DOI:** 10.1101/076224

**Authors:** Matthias Kaiser, Florian Jug, Olin Silander, Siddharth Deshpande, Thomas Pfohl, Thomas Julou, Gene Myers, Erik van Nimwegen

**Affiliations:** Biozentrum, University of Basel, and Swiss Institute of Bioinformatics, Basel, Switzerland; Max Planck Institute of Molecular Cell Biology and Genetics, Dresden, Germany; Department of Chemistry, University of Basel, Basel, Switzerland

## Abstract

Bacteria adapt to changes in their environment by regulating gene expression, often at the level of transcription. However, since the molecular processes underlying gene regulation are subject to thermodynamic and other stochastic fluctuations, gene expression is inherently noisy, and identical cells in a homogeneous environment can display highly heterogeneous expression levels. To study how stochasticity affects gene regulation at the single-cell level, it is crucial to be able to directly follow gene expression dynamics in single cells under changing environmental conditions. Recently developed microfluidic devices, used in combination with quantitative fluorescence time-lapse microscopy, represent a highly promising experimental approach, allowing tracking of lineages of single cells over long time-scales while simultaneously measuring their growth and gene expression. However, current devices do not allow controlled dynamical changes to the environmental conditions which are needed to study gene regulation. In addition, automated analysis of the imaging data from such devices is still highly challenging and no standard software is currently available. To address these challenges, we here present an integrated experimental and computational setup featuring, on the one hand, a new dual-input microfluidic chip which allows mixing and switching between two growth media and, on the other hand, a novel image analysis software which jointly optimizes segmentation and tracking of the cells and allows interactive user-guided fine-tuning of its results. To demonstrate the power of our approach, we study the lac operon regulation in *E. coli* cells grown in an environment that switches between glucose and lactose, and quantify stochastic lag times and memory at the single cell level.

## Introduction

Most bacterial species live in highly variable environments. To survive such environmental changes, bacteria must quickly adapt to the changing conditions, and they most commonly accomplish this by regulating gene expression [1, 2]. Since the molecular mechanisms that underlie gene expression are subject to thermal and other sources of noise, gene expression is inherently stochastic, and genetically identical cells in a homogeneous environment can exhibit significant variation and fluctuations in gene expression levels [3, 4]. Consequently, to gain a quantitative understanding of gene regulation and of the effects of stochasticity, it is necessary to study gene regulation in single cells.

Arguably the most common approach to studying gene expression at the single cell level is quantitative fluorescence time-lapse microscopy (QFTM) in which growth and gene expression of single cells are tracked within a microcolony of bacteria on an agar patch. Many exciting insights into the mechanisms and dynamics of gene expression have resulted from such studies, e.g. [5, 6]. However, these studies typically focus on single cell dynamics in a single environmental condition at a time. To gain deeper insights into regulation at the single cell level, it will be necessary to dynamically track how single cells *respond* in gene expression as they are exposed to *changing environmental conditions*. Although it is possible to expose cells to changing conditions on agar patches by growing the microcolonies between the coverslip and a membrane or an agarose gel over which different media are flown [7–9], observations are limited to short periods of time. After a few divisions the microcolonies grow out of the field of view or the cells start to form multi-layers.

Recently developed microfluidic devices solve these problems by growing cells in confined compartments and using flow to supply nutrients and wash away excess cells, allowing observations over long time scales [10–12]. An especially attractive design is the so called Mother Machine [10], in which cells grow in single-file within narrow side-channels connected to a main channel with nutrient flows. In this way, each growth channel (GC) provides a single independent lineage of growing and dividing cells that can be tracked over long time scales. Unfortunately, all the current designs expect a single media to be used as input. Consequently, growth conditions can only be altered by manually switching of the inputs, e.g. [13], which precludes growth conditions to be varied in a controlled manner. To be able to study gene regulatory dynamics at the single cell level, there is thus a strong need for a microfluidic setup that combines the advantages of previous devices with the ability to precisely and dynamically control the growth environment.

Another key challenge is the automated image analysis of the resulting time-lapse movies. Cells need to be automatically tracked, which consists of segmenting the images to identify the cells and linking these cells across images from neighboring time points. In addition, once the cells have been successfully tracked, additional methods are needed to quantify their size and the number of fluorescent molecules per cell. A number of analysis tools have become available for analyzing QFTM data from microcolonies [14–16]. First, the vast majority of implemented algorithms rely on fluorescence to segment the images, requiring one fluorescent channel to be used for identifying the cells. Moreover, these methods do, unfortunately, perform relatively poorly on time-lapse data from microfluidic devices of the type studied here because, being immersed in fluid, the cells can exhibit much larger movements between consecutive imaging frames than occurs with cells on agar patches. Today there is, in fact, no software tool available to track cells from microfluidic time-lapse movies. This is a significant impediment to the wider adoption of this exciting technology.

We here present an integrated experimental and computational setup that allows accurate quantitative study of gene regulation at the single-cell level in varying environments. In particular, we introduce the Dual Input Mother Machine (DIMM), a new dual-input microfluidic device that combines the Mother Machine design with precise environmental control. Further, we present the Mother Machine Analyzer (MoMA), a software tool that uses a new computational approach for joint segmentation and tracking, and provides highly novel ‘leveraged editing’ capabilities. That is, MoMA provides an interactive graphical interface that lets users browse through the current tracking solution, while displaying alternative segmentation and tracking solutions that can be selected through clicking on them. MoMA then automatically recalculates an optimal tracking incorporates these selections as additional constraints. In this way, corrections introduced by the user automatically percolate to neighboring time points, correcting multiple mistakes at once, leading to a dramatic increase in curation speed.

## Results

The design of our DIMM microfluidic device closely follows that of the original Mother Machine [10]. It consists of a main channel that carries nutrient-containing media and an array off small dead-end growth channels (GCs) that branch from the main channel (Fig. 1A+C). Nutrients diffuse from the main channel into the GCs in which cells are trapped (Fig 1C), and as the cells in the GCs grow and divide, cells closest to the GC exit are pushed out and are transported away by the flow in the main channel. In contrast to previous designs, our device has a dual input junction and mixing serpentines which allow for arbitrary time-dependent mixing of two input media. The two inputs meet in a Dial-A-Wave junction (DAW junction) [17] consisting of two inlets and three outlets (Fig. 1B). While the middle outlet feeds into the main channel of the device, the outer outlets function as waste channels and allow the flow in the middle outlet to vary from carrying only one of the two inputs (black in Fig. 1B), to carrying only the other input (green in Fig. 1B), without getting backflow into the inactive inlet. By varying the relative pressure on the two inputs using programmable syringe pumps, arbitrary time-dependent mixtures of the two input media can be realized, such as dynamically changing concentrations of particular nutrients or stressors.

As a proof of principle, we subjected *Escherichia coli* cells to media that were switching every four hours between glucose and lactose as carbon source (Fig 1). We used a modified MG1655 strain that carries a translational *lacZ-GFP* fusion at its native locus [6]. Time lapse movies of 22-24h were obtained in duplicatae for three different conditions (constant M9 + 0.2% glucose, constant M9 + 0.2% lactose, and switching between these two media every 4h), taking a frame every 3 minutes. An example movie of a 24 hour switching experiment is available [here](https://www.youtube.com/watch?v=2Tznm868fmc). To quantify the auto-fluorescence of the cells, we also exposed wild-type MG1655 cells to conditions switching between glucose (12h) and lactose (12h). We thus analysed 7 different time-lapse experiments all together, amounting to data from 180 GCs, more than 10’000 full cell cycles, and more than 500’000 single cell observations.

The analysis of the acquired image sequences is performed in two sequential steps; a preprocessing step, and the subsequent analysis by MoMA (*cf.* Fig. 1E). The preprocessing registers the frames of a movie sequence at sub-pixel accuracy to correct for jitter and stage drift. We then crop all GCs from this registered image sequence and store them on disk, resulting in one image sequence per GC. MoMA can load these stored GC image sequences for automated tracking and data analysis. In contrast to most image analysis tools, which first segment each of the images, and then try link the cells between images, MoMA jointly optimizes the segmentation and the linking in one multi-hypotheses assignment tracking algorithm. To make this joint optimization tractable, MoMA casts this tracking problem as an integer linear programming (ILP) problem. Importantly, the tracking can be performed on phase contrast images, leaving all fluorescent channels for measurement of gene expression, and allowing for tracking of non-fluorescent (e.g. wild-type) cells. For each phase contrast image, MoMA finds an overcomplete hierarchy of segmentation hypotheses, i.e. potential cell segmentations. For each set of such segmentation hypotheses MoMA enumerates potential ‘fates’, called *assignments*, to link the cells across time points, i.e. all possible cell movements, cell divisions, and GC exit events. A cost is assigned jointly to all segmentation hypotheses and assignments. Low costs can be interpreted as a high likelihood of the associated segmentation/assignment for explaining the image sequence. Additionally, MoMA implements constraints that describe how hypotheses and assignments must depend on each other in order to assemble sound tracking solutions. For example, cells cannot move passed each other, only the top cells can exit the GC, and so on. MoMA then uses Gurobi, a potent off-the-shelve solver for mixed linear integer programs, to find the tracking of lowest overall cost. This is equivalent to finding the maximum a posteriori solution in Bayesian statistics.

In practice, a tracking is computed automatically when MoMA starts and the image sequence of a preprocessed GC is loaded. An interactive graphical user interface then allows users to browse the possible solutions, identify potential errors, and if necessary, correct them. In contrast to existing methods, where data curation is performed by explicitly editing the segmentations or linking graphs, MoMA offers several different ways for browsing through alternative segmentation and assignment hypotheses, and for selecting among these. Once a user made an adjustment, e.g. by selecting an alternative segment or link, MoMA formulates the user’s choice as an additional constraint and restarts the optimization of the underlying ILP in order to find the new maximum *a posteriori* solution incorporating this constraint. In this way corrections automatically percolate to nearby time points. A tutorial video showing leveraged editing in MoMA is available on Github at https://github.com/fjug/MoMA/wiki/MoMA%20Movies. After the user gets to know the user interface, it typically takes only 1 to 2 minutes to fully curate 100 frames (see Fig. 1F), corresponding to 5 hours of real time with our settings. Once curation is completed, the correct tracking can then be exported (as shown in the supplementary movie at https://github.com/fjug/MoMA/wiki/MoMA%20Movies). The exported file contains information about the sizes and fluorescence levels of each cell across all time points in its cell cycle, the locations of the cells in the GC, each cell’s parental lineage, etcetera.

**Figure 1:**
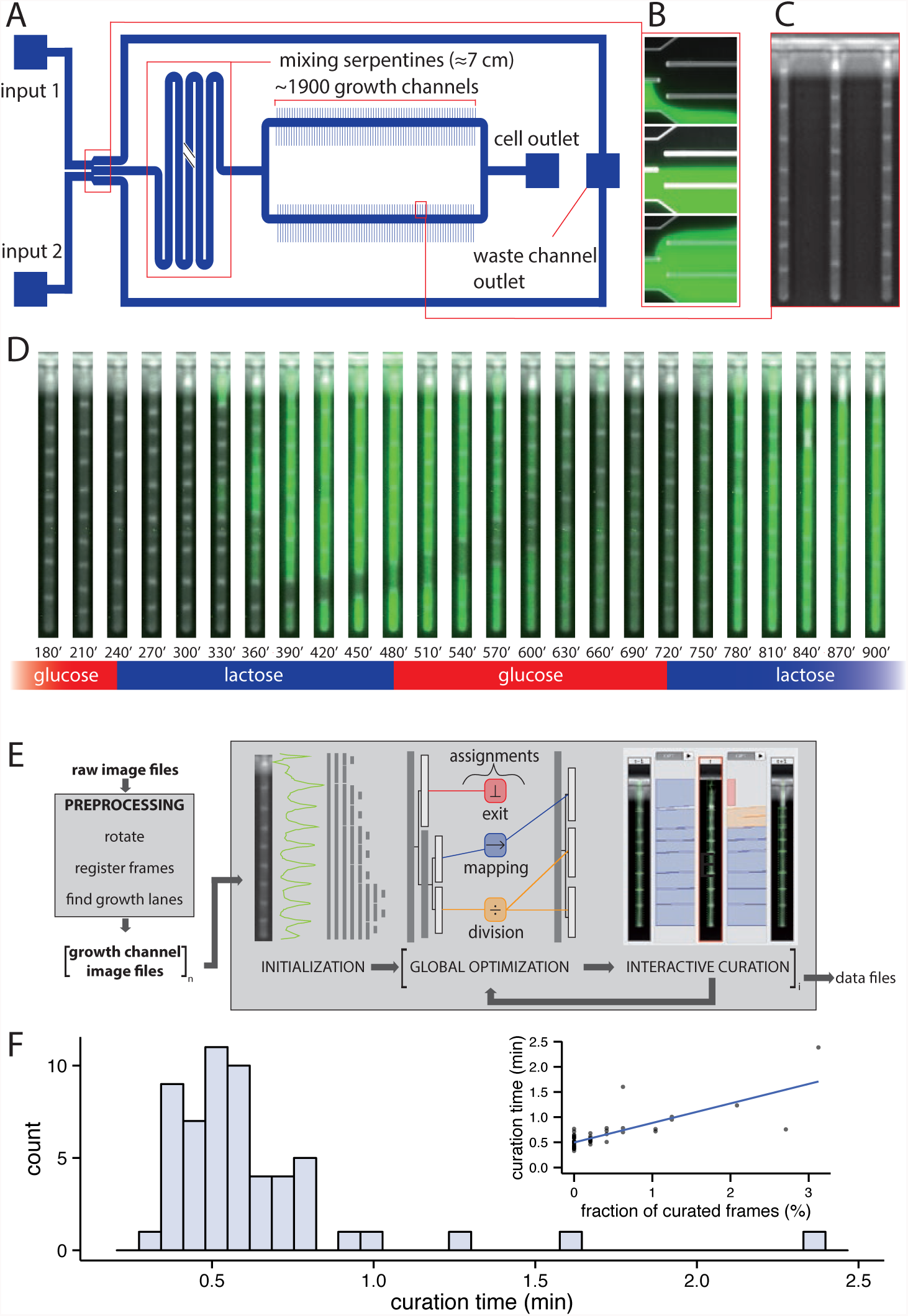
A) Overview of the Dual Input Mother Machine (DIMM) design, B) DAW-junction in three different switching states, top: 100% from input 1 (unlabeled) and 0% from input 2 (green), middle: 50% from input 1 and 50% from input 2, bottom: 0% from input 1 and 100% from input 2, C) Phase contrast image of *E. coli* cells in three GCs during growth in the DIMM, D) A time series of a single GC containing *E. coli* cells expressing *lacZ-GFP* from the *lac* promoter while being exposed to alternating switches between media containing glucose and lactose as a carbon source, E) Overview of the different steps in the MoMA analysis pipeline, F) Expenditure of time for curating mother machine data using MoMA, main: Distribution of curation times per 100 frames (*∼*30 frames per cell cycle), inset: curation time per 100 frames against what fraction of frames needed curation.

MoMA and its documentation including tutorial videos are available on Github at https://github.com/fjug/MoMA. There we also provide downstream analysis tools that we used to convert the raw cell size and fluorescence outputs from MoMA into physically meaningful quantities. In particular, we refine the size estimates using pixel intensity profiles across each cell (Suppl. Methods and Fig. 7). We find that cell size across the cell cycle is very well described by an exponential growth curve, and much better than by a linear growth curve (Fig. 8).

Second, we developed a novel procedure for estimating the total fluorescence of each cell, which significantly improves over simply summing intensities across pixels within the cell. In particular, we have found that the fluorescence of each cell spreads significantly beyond the cell outline, and falls in an approximately Lorentzian manner as a function of distance to the cell. As a consequence, background fluorescence levels tend to increase as a function of the fluorescence of neighboring cells (Fig. 9) and we use a Lorentzian mixture model to accurately estimate the fluorescence of a given cell (Suppl. methods). To correct for the cell’s auto-fluorescence, we performed an independent experiment with wild-type MG1655 (i.e. without the *gfp* gene) and found that auto-fluorescence is proportional to cell volume (Fig. 10). We estimated the auto-fluorescence per unit volume from this data, and used this to subtract auto-fluorescence from the data obtained with the *lacZ-GFP* strain.

Finally, in order to quantify the number of fluorescent molecules per cell, we needed to estimate the conversion factor between measured fluorescent intensity and absolute number of GFP molecules. Whenever a cell divides, the fluorescence of the mother is divided approximately equally to the two daughter cells, and Rosenfeld et al. [18] previously developed a procedure for estimating the conversion factor by assuming fluctuations in the fluorescence of the two newborn siblings is entirely due to the random binomial distribution of molecules between them. Here we developed a adaptation of this method that significantly improves on the method of Rosenfeld et al. [18] in several ways. First, in order to avoid the confounding effects of GFP synthesis, the analysis is best performed in a condition where GFP is expressed but no GFP synthesis occurs. Data from the glucose phases of the experiments with switching conditions (see Fig. 2A) precisely fit this requirement, and provide much more extensive data than were available in [18]. Second, we accounted for the slow decrease of fluorescence during the glucose phase due to a combination of protein decay and bleaching of the GFP by estimating this fluorescence decay and correcting for it to estimate GFP levels of the newborn siblings at division. Third, our analysis of the fluorescence levels of newborn sibling pairs showed that, that in addition to the expected binomial fluctuations, there are significant fluctuations in the sizes of the new siblings. Moreover, the fluorescence fluctuations correlate with these size fluctuations (Fig. 12). To account for this, we developed a new Bayesian procedure that takes both the binomial and size fluctuations into account and rigorously calculates the posterior distribution for the conversion factor between GFP fluorescence and the number of *lacZ-GFP* tetramers (Suppl. Methods and Fig. 13). With this procedure we find that size fluctuations in fact dominate the total fluctuations at division and that, when growing in lactose, cells contain about 3000 − 6000 GFP molecules at birth and 6000 − 12000 GFP molecules just before division.

**Figure 2:**
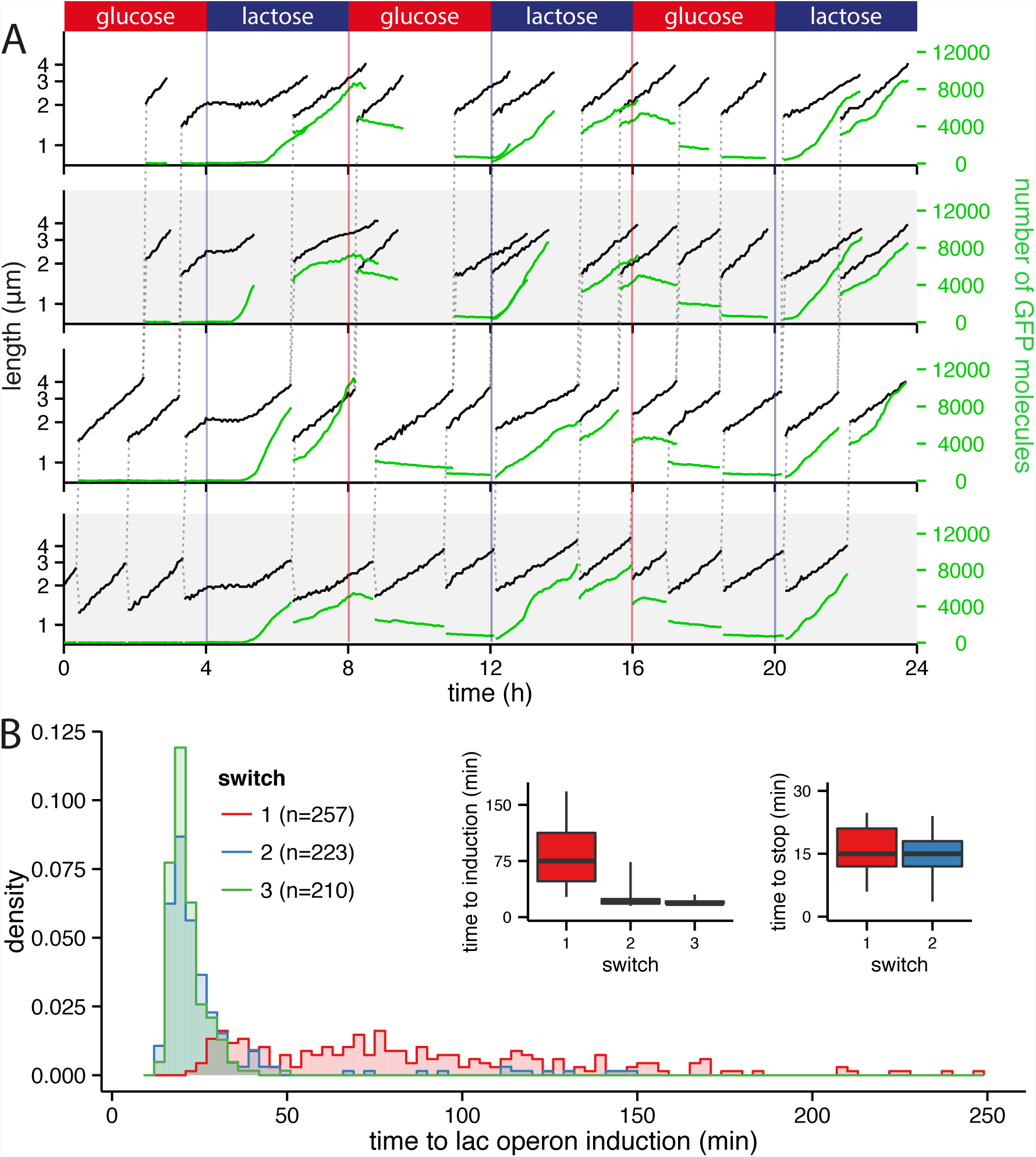
A) Dynamics of growth and gene expression in lineages of single cells in an environment that switches between M9 + 0.2% glucose and M9 + 0.2% lactose every 4 hours. Cell size (black, logarithmic scale) and expression of *lacZ-GFP* (green, linear scale) are shown as a function of time for a lineage of cells at the bottom of the GC (bottom row) together with first and second generation offspring cells (top three rows). The dashed vertical lines show the lineage of cell divisions by connecting each mother cell to its two daughter cells. B) Distribution of lag times to *lacZ-GFP* induction in single cells at the first (red), second (blue) and third (green) switch from glucose to lactose. The insets show boxplots (median, interquartile range, 5 and 95 percentiles) of the lag-time distributions (left) and the distribution of waiting times until termination of *lacZ-GFP* production (right).

Figure 2 illustrates the kind of detailed quantitative information regarding gene regulation at the single-cell level that our methodology provides. During the initial glucose phase cells grow exponentially while *lacZ* expression is close to zero. As soon as the medium switches to lactose, essentially all cells arrest their growth immediately, i.e. within the 3 minutes between measurements. During this growth arrest *lacZ* expression remains low until the lac operon is induced and fluorescence starts increasing rapidly. The length of this lag phase to induction of the *lac* operon fluctuates significantly from cell to cell, and some cells do not induce at all within the 4 hour lactose phase. Interestingly, the time at which *lacZ* expression is first observed coincides almost precisely with the restart of exponential growth. Moreover, instead of growth restarting slowly and increasing in speed as lacZ levels increase, it appears cells grow at essentially ‘full speed’ immediately after restart. When the condition switches back to glucose there appears to be little effect on the exponential growth of the cells, but we observe a cessation of *lacZ-GFP* production shortly after this switch. In contrast to the lag times to *lacZ-GFP* induction, the times until cessation of *lacZ-GFP* expression are very narrowly distributed around 15 minutes, which may be dominated by the folding time of the GFP.

At the end of the second glucose phase the GFP levels have been diluted almost to background levels, but no growth arrest is observed at the second switch to lactose, and *lacZ-GFP* expression is immediately induced for cells that previously induced *lacZ-GFP*. This suggests that the proteins which remain from the lac-operon expression during the previous growth in lactose act as a ‘memory’, confirming previous observations [12]. To quantify some of these qualitative observations, Fig. 2B shows the distribution of the single-cell lag times to *lacZ* induction for the first (red), second (green), and third (blue) switch to lactose. There is a very wide distribution of lags in the first switch, with lags over an hour for many cells, and lags exceeding the entire 4 hour lactose phase for some cells. In the second switch the large majority of cells show lags of less than 30 minutes, although some cells with long lags remain. No more long lags are observed for the third switch. The fact that growth restarts at essentially ‘full speed’ as soon as *lacZ-GFP* synthesis is detectable, i.e. when *lacZ* levels are still much below steady-state levels, is especially intriguing. If the restart of growth depends directly on carbon source metabolism, than this observation would suggest that only low levels of *lacZ* are needed for optimal growth. This would then raise the question as to why steady-state *lacZ* levels are so much higher. Alternatively, it may be that growth-rate is not directly limited by the carbon source. In this case the coincidence of growth restart and *lacZ-GFP* induction may indicate that these events are coupled through a common sensory mechanism. These hypotheses are interesting topics for further study.

## Conclusions

We have here presented an integrated experimental and computational setup for quantifying gene expression dynamics at the single cell level over long periods of time in dynamically changing environments that are precisely controlled. This methodology opens up a wide array of possibilities for studying gene regulatory mechanisms at the single cell level. As an illustration, we here measured lag times in single cells upon switches between different carbon sources, but many other types of investigations would be possible. For example, by varying the time between switches, the length distribution of memory effects in single cells could be determined. Our setup can also be generally used to quantify how expression fluctuations affect fitness, i.e. growth rate, at the single cell level. Other possibilities include measuring how the distribution of gene regulatory responses across single cells depends on the concentration of an inducing nutrient or stress. Alternatively, by using short pulses or oscillating levels of inducers, we can measure how single cell gene regulatory responses depend on the time dynamics of the inducer, which is potentially highly informative for investigating regulatory network architecture. In summary, we believe that the integrated experimental and computational methodology that we present here will be an important tool for studying gene regulatory mechanisms at the single cell level.

## 1 Experimental methods

### 1.1 Design and fabrication of the microfluidics devices

E. coli cells take on different sizes depending on the media they are grown in, e.g. LB versus M9 minimal medium. Since the GCs aim to trap the cells growing in single file, the width of the channels needs to match the width of the cells as closely as possible. To account for this, our DIMM device has contains channels with a range of widths. For the results presented here, the GC sections were *∼*0.8µm x *∼*0.8µ. These dimensions worked nicely with cells growing in M9 + 0.2% glucose or 0.2% lactose respectively. Experiments with other media and strains might require slightly different dimensions. In order to reduce the flow rates compared to the original mother machine device the dimensions of the main channels were reduced to a diameter of 6µm x 50µm in the device presented here. The resulting flow rates are discussed in more detail in the section discussing loading and flow control. The device was designed using the Software AutoCAD^®^ from AUTODESK^®^. These designs are available upon request from the authors. We outsourced both the production of the photomask and the production of the masters to pour the PDMS devices from. A 5" quartz mask with chrome layer was ordered from the Compugraphics Jena GmbH. Using this mask micro, resist technology in Berlin produced the master using UV-lithography with SU-8 photoresists (for more details see [10]). To make the chips, we use the Sylgard Elastomer Kit 184 with a 1:10 curing agent to base ratio. Curing was performed at 65°C overnight or longer. Harris Uni-Core 0.75mm biopsy punches were used to create in-and outlets. Before bonding, both the PDMS mold and a cover slip were washed with isopropanol and dried with air. Surface activation was done in a plasma cleaner (PDC-32G, Harrick Plasma) operated at high intensity with vacuum at 1500 mTorr for 40 seconds. After bonding the devices were incubated at 65°C for at least 1h.

### 1.2 Performance of the environmental control

The design presented here not only allows switching between different media but also allows for continuous control over the ratios of two different input media. Because flows in micro-channels are strictly laminar only diffusive mixing occurs at these scales [17]. To keep the design simple we introduced 2D mixing serpentines to the device. These serpentines guarantee that the media coming together in the DAW junction are flowing together long enough to allow for diffusive mixing before the mix reaches the cells. The required length of these mixing serpentines depends on the flow speed (fluid velocity), the width of the micro-channels and the diffusion coefficient of the molecule of interest in the medium used [19]. To demonstrate mixing in the device we used M9 minimal medium labeled with fluorescein (1µg/ml) mixed with unlabeled M9 minimal medium. Figure 3A demonstrates how the system presented here can generate different mixing ratios and thus can be used to precisely control the growth environment. As the relative flow rates of the two inputs are varied, the fluoresence measured downstream in the main channel varies linearly when the relative flow rates are in the regime 2/3 to 3/2. Figure 3B shows the normalized fluorescent intensity along a section through the main channel dowstream of the mixing serpentines at different flow regimes. Because of small imperfections in the mold the intensity profile is not perfectly symmetrical even in the unmixed state (black line). However in the different mixed states, the shape of the profile stays the same indicating complete mixing is guaranteed in the flow regimes tested here.

### 1.3 Strains and growth conditions

Strains were kept at −80°C in LB+15% glycerol stocks and were streaked on LB plates before experiments. M9 minimal medium supplemented with 0.2% glucose or 0.2% lactose was used for precultures and microscopy experiments. The night before an experiment an overnight culture was inoculated from a single colony in minimal medium with the same carbon source that the cells were to experience in the initial phase of the experiment. The next day, cells were diluted 1/100 into 5-10ml of fresh medium with the same carbon source. Cells were harvested after 4-6h to be concentrated and loaded into the microfluidic device (typically, a culture at OD*≈* 0.2 is concentrated 100×). Growth temperature was 37°C for both precultures and the experiments.

### 1.4 Priming and loading of microfluics devices

The DIMM design presented here has two inlets and two outlets. This leads to some complications in the cell loading process compared to the original Mother Machine design. Here we describe the adjusted loading procedure we developed. As described in [10], a mixture of salmon sperm DNA (10mg/ml) and BSA (10mg/ml) (mixing ratio 1:3) is used to passivate the device before loading the cells. The salmon sperm DNA is denatured at 95°C for 10 min and is mixed with the BSA after cooling down. This passivation buffer is also added to the growth medium in the experiment in a concentration of 1/100. In addition, one medium was always labeled with non-fluorescent microspheres (Polybead^®^ Polystyrene 1*µ*m Beads) to monitor medium flow at the DAW junction. As shown in Figure 4 the two DAW waste channels cannot be pressured separately because they both end in the same outlet. Therefore to prevent blockage of one of the waste channels by passivation buffer it is recommended to flow water into the waste channel outlet while the passivation buffer is loaded into the cell outlet. Once the main channel (with the GCs) is filled with passivation buffer and the inlets (input 1 and input 2) are full of liquid (mixture of water and passivation buffer), both the flow of water and of passivation buffer can be stopped. The device is now incubated for ca. 1h at 37°C before the loading of the cells is started.

**Figure 3:**
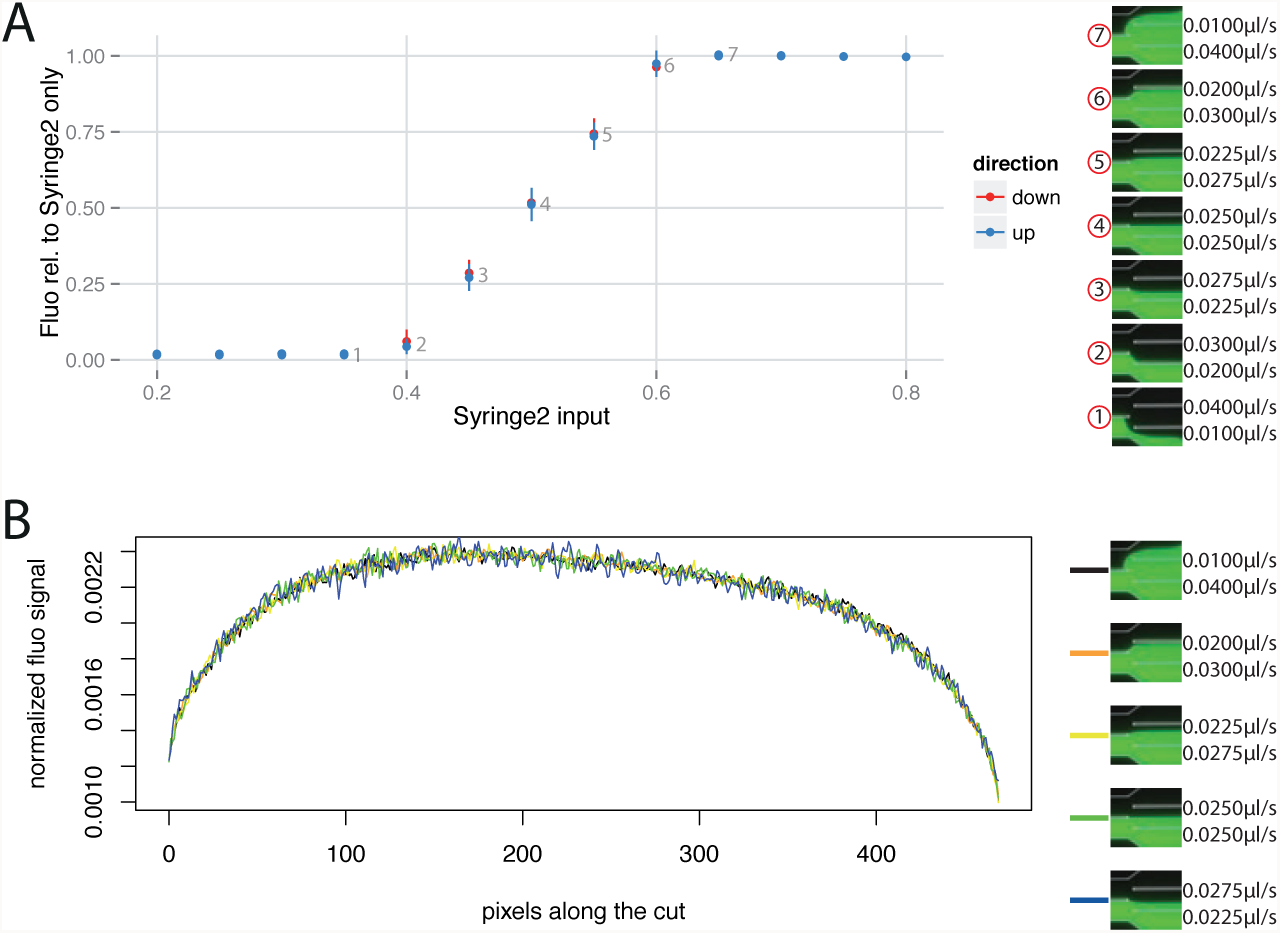
Mixing of fluorescein labeled medium with nonlabeled medium at different input flow rate ratios. A) Total fluorescence was measured in a square region in the main channel downstream of the mixing serpentines as the input ratio was changed in a stepwise manner from 0% fluorescein input to 100% fluorescein input (up: blue) and back to 0% fluorescein input again (down: red). The fluorescence measured for the mixture relative to the fluorescence measured for pure fluorescein labeled medium (Syringe 2 only) is plot against the input of Syringe 2 as a fraction of the total input from both Syringe 1 and 2. B) Normalized fluorescence along a section through the main channel downstream of the mixing serpentines at different mixing ratios.

**Table 1:** strains used in the study.

| Strain Name | Relevant Genotype | Ancestor | Source or Reference |
| --- | --- | --- | --- |
| MG1655 | *E. coli* K12 wild-type |  | [20] |
| ASC662 | *lacZ-GFP* | MG1655 | [6] |

**Figure 4:**
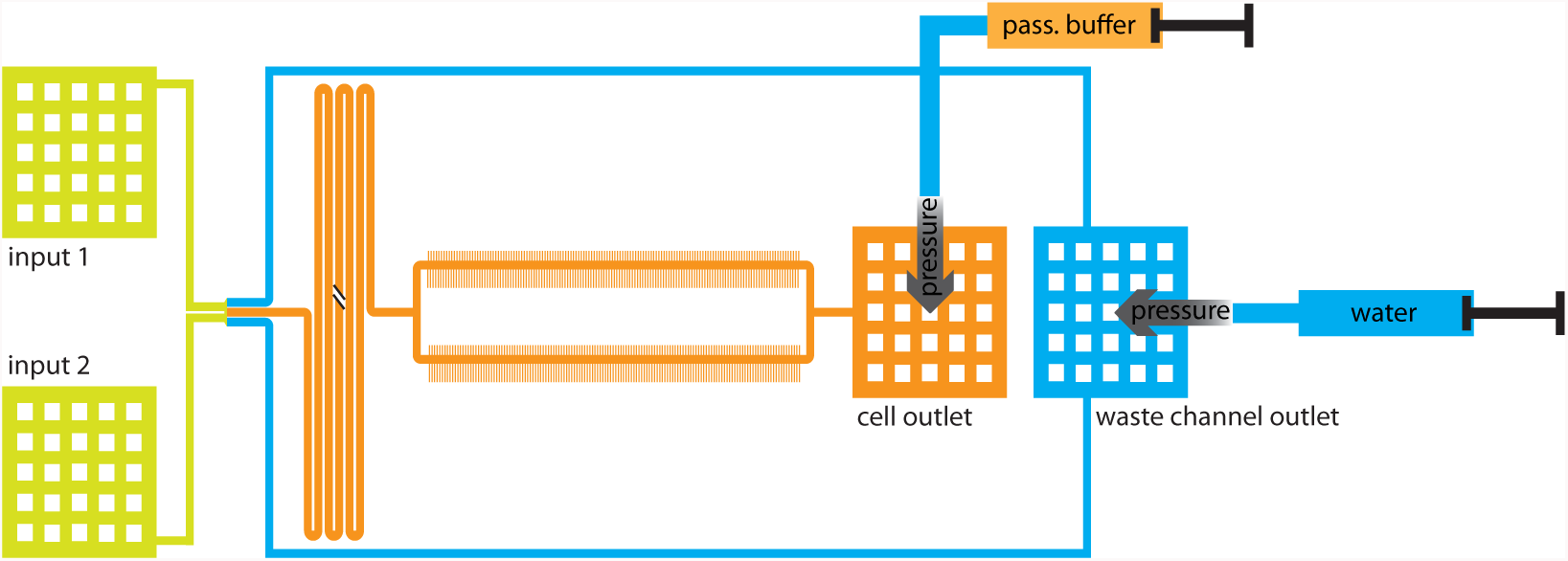
Loading of the passivation buffer into the main channel: To prevent blocking of the waste channels by passivation buffer, the waste channels are loaded with water through the waste channel outlet (blue) while loading of passivation buffer is done through the cell outlet (orange). Putting both outlets under pressure assures complete loading of the main channel with passivation buffer while the waste channels stay clear of passivation buffer.

After the passivation step cell loading can begin. To get rid of the passivation buffer, the two inlets are connected to the pumps with the two different media that will be used in the experiment. At this point the tubing for the waste outlet can also be installed and connected to a waste container. Both pumps are now set to a flow rate of 0.025µl/s. When all channels are clear, this flow regime will lead to a 50:50 ratio between the two inputs at the DAW junction. If the device leaks at this point or fails to establish a 50:50 ratio at the DAW junction (one medium is labeled with beads to check the flow under the microscope), most likely the resistance of some channel is altered by a blockage and the device cannot be used. If the device works properly, the DAW junction can be switched to the medium that will be used first. This step is necessary to ensure that the cells that are loaded afterwards only encounter the media condition in which they will begin growth. For a complete switch we use flow rates of 0.01µl/s for the inactive inlet and 0.04µl/s for the active one (Figure 3A). After a few minutes (depending on the flow rates) the main channel and cell outlet should be free of the medium from the initial input and the cell loading process can begin. The cells are harvested in exponential phase and are concentrated by centrifugation (*∼* 100-200x). Once the device is fully switched to the desired input, one can load the cells using a 1ml syringe into the tubing that will later serve as the waste tube. This tubing is inserted into the cell outlet and can be pressured by hand to flow the cells into the main channel (the loading process is observed under the microscope). It is important not to stop the flow at the inlets during the whole loading process. This allows cell loading without getting cells into the inlets where they might become stuck and grow. Once the cells reach the GCs we used a custom made clamp to hold a precise level of pressure on the 1ml syringe for cell loading. The pressure here has to be continuously adjusted to make sure the cells stop flowing in the main channel and can enter the GCs. As shown in Figure 5 there is a constant flow through the inlets and the waste channels (green) while the main channel is pressured to achieve zero flow where the GCs are (red). If cells move up to the DAW junction they are removed through the waste channels and the inlets stay clear. Loading continues until a satisfactory number of channels contain cells (typically 20 to 60 minutes). When complete, the 1ml syringe used for loading is removed, and the end of its tubing is put into the waste container together with the tubing from the waste outlet. After loading the cells are allowed to recover for at least 2h before the experiment starts.

Growth media are delivered from air-tight glass syringes (Hamilton) that are connected to the device using PTFE tubing with an inner diameter of 0.56mm and an outer diameter of 1.07mm. The syringes are controlled by two low pressure pumps (Cetoni GmbH) so that the total flow during the experiment is 0.05µl/s.

**Figure 5:**
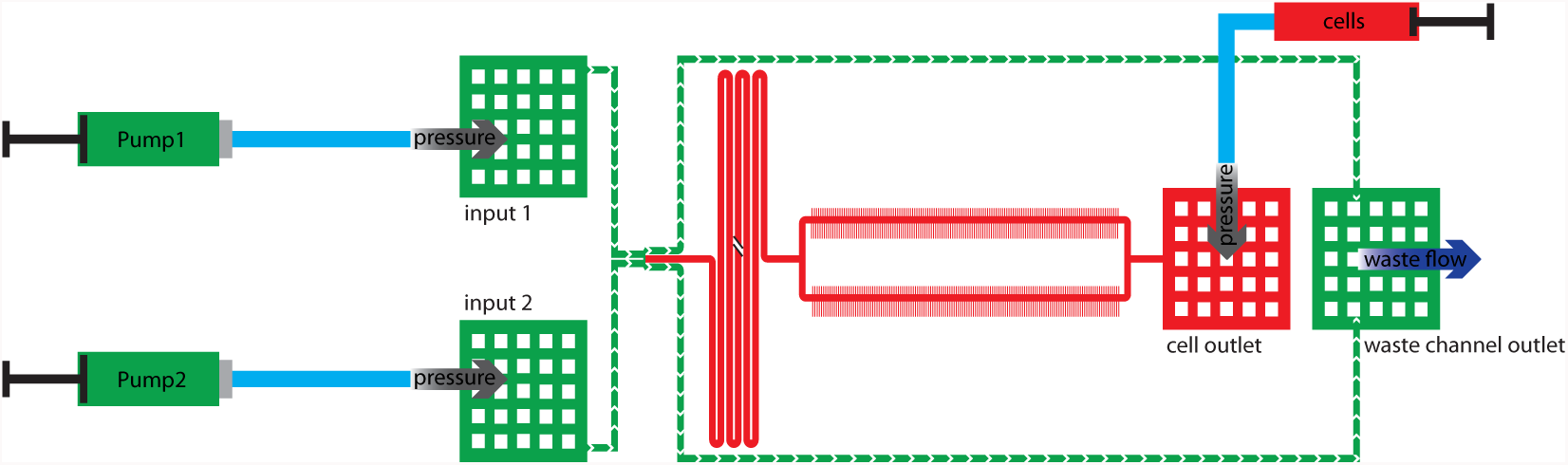
Cell loading procedure: A constant flow in both inlets (input1 and input2) prevents cells entering the inlets during the loading. The concentrated cell solution can be loaded through the cell outlet. First some pressure is applied to fill the whole main channel with cells. Afterwards the pressure is controlled to maintain zero flow in the main channel (red) while there is a constant flow through the inlets and in the waste channels (green) to remove cells that make it up to the DAW junction.

### 1.5 Microscopy

An inverted Nikon TI-E microscope equipped with a motorized xy-stage with linear encoders was used to perform the experiments. All experiments were performed in an incubator maintained at 37°C. The sample was fixed on the stage using metal clamps and focus was maintained using the Perfect Focus System from Nikon. Images were recorded using a CFI Plan Apochromat Lambda DM 100x objective (NA 1.45, WD 0.13mm) and a CMOS camera (Hamamatsu Orca-Flash 4.0). The setup was controlled using Micro-Manager [21] and timelapse movies were recorded with its Multi-Dimensional Acquisition engine (customized using *runnables*). Every 3 minutes one phase contrast image and one GFP fluorescence image were acquired, typically for 6 different positions. Phase contrast images were acquired using 100ms exposure with the transmitted light source at full power (CoolLED pE-100). Images of GFP fluorescence were acquired using 2s exposure, illuminating the sample with a Lumencor SpectraX (Cyan LED) set to 17% and dimmed using a ND4 filter in the light path; the excitation (475/35nm) and emission filters (525/50nm) were used with a dichroic beam-splitter at 495nm. For the switching experiments images of the DAW junction were also acquired. Here the GFP channel was replaced with an additional phase contrast image with a short exposure time (10ms) to visualize the beads in the flow.

### 1.6 Preprocessing of microscopy data

MoMA expects to be given image datasets in which a single GC is present, with the GC opening at the top, and with phase contrast being the first channel. With our microfluidics design, the camera field of view covers ca. 30 GCs so the images must be split into individual GCs and preprocessed in order to match MoMA’s requirements. The preprocessing consists of the following tasks:

1. Load the microscopy dataset, one position at a time, in a format-independent manner using the Bio-Formats library (in order to open a specific position, one must use the Java API rather than functions available in ImageJ).
2. register all frames to the first frame of the first channel in order to correct the sample drift over time, as well as the jitter introduced by acquiring multiple positions in parallel. To do this, we develop HyperStackReg, a custom extension of the StackReg ImageJ plugin that is able to handle hyperstacks i.e. datasets with several channels.
3. Crop the image to keep only the area of the GCs and rotate the images (so that GC opening is at the top).
4. Save images as a tiff dataset with one file per frame.
5. Straighten the image so that GCs are oriented vertically (using bicubic interpolation).
6. Identify the GCs in the first phase contrast frame and save one dataset per cropped GC.

Steps 1. to 4. are run in Fiji while steps 5. and 6. are done by mmpreprocess, a custom standalone Java program. Because of ImageJ1 limitations in handling multidimensional stacks, mmpreprocess and MoMA only handles datasets with one file per frame. This is likely to change in the future and will be documented on MoMA’s wiki (along with the updated preprocessing scripts). Also, in order to preprocess datasets from the command line, fiji must be run using a virtual window environment (using Xvfb), since the headless mode is not compatible with some important ImageJ features.

We release the HyperStackReg ImageJ plugin as well as the mmpreprocess executable on MoMA’s wiki. We also provide a bash script that allows preprocessing of all images acquired at one position in a single step, along with the necessary ImageJ macro. This is meant as a starting point for users to write a preprocessing workflow tailored to their needs. In particular, loading different image formats with Bio-Formats is likely to require adjustments.

## 2 The *Mo*ther *M*achine *A*nalyzer, MoMA

Today’s predominant tracking methods originated in the 1960s [22, 23] and were developed to track single or a hand-full of objects with complex distinguishing features such as ships or airplanes. However, here we require the tracking of objects that are visually almost identical. In some cases this can be resolved by maintaining multiple association hypotheses over multiple time-points [24]. However, although particle trackers and state space models can produce high quality results, proofreading (data curation) is always required in order to guarantee error-free tracks. Notably, computer-assisted approaches for proofreading are usually not related to the automated method that produced the automated results in the first place.

Interactive error correction is rarely part of available tracking systems and usually turns out to be difficult to implement and integrate, leaving the user with an inflexible patchwork of multiple tools. Part of the reason for this is the way classical tracking models work. Their local and iterative solvers are highly specialized, not offering intrinsic possibilities to constrain the space of possible solutions in a a user-driven way. In other words, they intrinsically do not offer any interaction capabilities that can be employed by users to prevent the tracking system from making certain mistakes.

*Assignment Models* promise to make a difference in all these respects. The novelty of this type of tracking system is the way in which solutions are found. A tracking problem is formulated as a global optimization problem under constraints that can be solved using discrete optimization methods. MoMA is based on such an optimization based assignment model which allows the user to furnish constraints in an interactive manner. Hence, we can offer unprecedented user interactions for data curation – a process we call *leveraged editing*.

In particular, MoMA offers the following leveraged editing primitives: (*i*) Forcing solutions to contain a selected cell (segment), (*ii*) forcing solutions not to include specific segments, (*iii*) forcing a cell to a given movement or division (assignment), or to (*iv*) avoid such an assignment, and (*v*) specifying the number of cells visible at a given time. We will show that the very nature of the underlying optimization problem allows us to seamlessly incorporate these leveraged editing primitives.

### 2.1 Automated Tracking

Here we briefly review the class of tracking methods called *assignment models* [25–29]. We provide sufficient technical detail to prepare the reader for Section 2.2, where we introduce the leveraged editing primitives used in MoMA.

#### 2.1.1 Tracking by Assignment

Tracking consists of two equally important tasks: Cells need to be segmented in each frame, and segments of the same cell in consecutive frames need to be linked. Tracking by assignment approaches these tasks by formulating and solving a joint global optimization problem. In this context, the segmentation task consists of selecting a subset of segments in each image, i.e. corresponding to the cells in the image. To do this the algorithm first generates a large collection of possible segment hypotheses that are contained in a (possibly heavy) oversegmentation of the images. Joint segmentation and tracking then boils down to enumerating many potential subsets of segments together with potential ways of linking (*assigning*) these between consecutive frames. To identify, among all these possible joint segment/assignment subsets, an optimal solution, each of the potential segments and assignments is given a cost. The costs of joint segment/assignment hypothesis aim to reflect how unlikely it is that the corresponding dynamics occurs in the real system, i.e. the corresponding movement, growth, and division of the cells in our system. That is, the total cost can be thought of as a negative log-likelihood of the segment/assignment hypothesis [26, 30] and the optimal solution minimizes the summed costs.

The cost function is designed to reflect the knowledge of domain experts. To give an example, in our application, the cost function for a cell division assignment that links one segment to two segments in the next frame contains a term that evaluates the size of the three segments to be linked which implements the physical constraint that the sum of the sizes of the two daughter cells should be similar and at least as large as that of the mother cell. Structural knowledge about which assignments can be chosen simultaneously is encoded in terms of constraints that ensure that only physically meaningful solutions can be chosen. That is, solutions that do not describe impossible events like cells popping into existence out of nowhere, cells moving to two places at once, etcetera. In our implementation, these constraints force or prohibit certain segments and assignments to be jointly contained in a segment/assignment solution.

Once the segmentation and tracking problem has been formalized in this manner in terms of costs and constraints, well-established discrete optimization methods can be used to obtain a solution that is (*i*) feasible, i.e., free of conflicts, and (*ii*) cost-minimal. In the following we will put these notions on formal grounds. A more in-depth description can be found in [29], where we described in great detail the assignment model upon which MoMA is based. In the next section we will briefly summarize this model in order to lay the foundation to understand the leveraged editing primitives we will introduce in Section 2.2.

#### 2.1.2 The Assignment Tracker within MoMA

First, an excess of segment hypotheses *H*^(*t*)^ is generated for each frame *t*, with many hypotheses partially overlapping and thereby providing alternative and mutually exclusive interpretations of where the cells are appearing in the image [29]. To represent possible solutions, a binary *segmentation variable h*^(*t*)^ is associated to each possible segment hypothesis in *H*^(*t*)^. Whenever *h*^(*t*)^ = 1, it indicates that this segment hypothesis is part of the proposed solution. Similarly, a set of assignment hypotheses *A*^(*t*)^ and associated binary *assignment variables a*^(*t*)^ are generated, that link segment hypotheses at time-point *t* to segment hypotheses at *t* + 1. That is, a mapping assignment 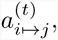 for example, connects two segment hypotheses 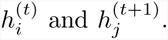. Again, whenever an assignment variable *a*^(*t*)^ = 1, this indicates that this assignment is part of the proposed solution.

Thus, a proposed segment/assignment solution consists of a selection of binary segmentation and assignment variables *v* that are set to 1. As mentioned above, a cost function is defined that associates to every such variable *v*, a cost *c*_*v*_ ∈ ℝ of including it in a solution. For details on the cost function used for the mother machine we refer to [29]. In a nutshell, the cost measures how much a segment/assignment deviates from the expected appearance/dynamic behavior of a bacteria cell. ‘qThe total cost *C* of a particular solution is then simply the summed cost over all active variables

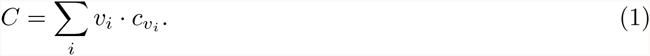

Linear constraints are used to restrict the solutions space to only include conflict-free and structurally sound solutions. As a simple example, two segment hypotheses which offer conflicting explanations of a particular pixel cannot simultaneously be active in any *feasible* solution. To introduce some notation that will be required below, we look in detail at one particular constraint. *Continuity constraints* ensure that each active segment at frame *t* (i.e. each cell) must be involved in exactly one assignment entering from frame *t −* 1 and must also be involved in exactly one assignment towards *t* + 1. In other words, each cell must have a past and a future. Formally, this statement can be written as

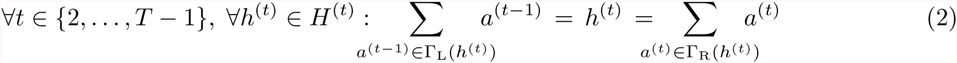

Here we image time frames ordered from left to right and use the notation Γ_L_(*h*) to denote the set of assignments directly to the left of segmentation variable *h* (i.e. its *left neighborhood*) and Γ_R_(*h*) to denote the set of assignments directly to the right of *h* (its *right neighborhood*). That is, the left neighborhood Γ_L_(*h*) is the set of all assignments entering *h* from the previous frame and the right neighborhood Γ_R_(*h*) is the set of all assignments leaving *h* towards the next frame. The equation above then says that, for each cell at time *t*, there should be one assignment in the previous time frame, and one in the following time frame.

#### 2.1.3 Finding the Optimal Solution

In tracking by assignment the globally optimal solution for picking a set of conflict free assignments of minimal cost is often found by solving an integer linear program (ILP) [26–29]. An ILP is an optimization problem that is fully specified by (*i*) an *objective function* that is a linear function of a set of variables *V*, and (*ii*) a set of *constraints* that are formalized as (in-)equalities on these variables. The space of *feasible* solutions is defined by all variable assignments that obey all constraints. An *optimal* solution is a feasible solution that minimizes the objective function.

The joint segmentation and tracking formulation introduced above is already in ILP form: The set of variables *V* comprises binary segmentation and assignment variables. The objective to minimize is the cost *C* defined in Equation (1). Note, that this is obviously a linear function of the variables *v* ∈ *V*. In Equation (2) we also gave an example of how constraints can be formalized as linear equalities.

Integer linear programming is a well-understood problem [31], and given the above formulation we can turn to off-the-shelf solvers like Gurobi™ to find an optimal solution. Although finding an optimal ILP solution is NP-hard, recent success solving relatively large tracking problems [26–29] suggests that assignment models pose well-natured instances to be solved as ILPs.

In the following we will make use of a particular feature of many ILP solvers, namely the ability to perform “warm-starts”. One speaks about a warm start if a solver can benefit from residual intermediate results created during a preceding optimization. This can speed-up optimization significantly as shown in [29].

#### 2.1.4 Reducing Redundancy by Substituting Segment Variables

The set of variables *V* contains variables *h*, for available segments, and *a*, for available assignments. However, note that whenever the segmentation variable for a segment *i* is active, i.e. *h*_*i*_ = 1, then at least one assignment *a* that involves a segment *i* must be active as well. Using these constraints, the segmentation variables can be removed from the model entirely [26–29]. That is, after adequate constraints are added to the ILP, each occurrence of 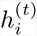 can be substituted by a sum over all assignment variables in 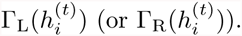.

### 2.2 Leveraged Editing

In this section we discuss how MoMA modifies the underlying optimization problem in response to user feedback. MoMA first of all provides the user with a graphical interface that allows the user to browse through the tracking solution that the optimization has provided for a given movie. The basic idea of leveraged editing is simple: When a user identifies a segmentation or tracking error, (s)he suggests the correct alternative or simply points at the error in the graphical interface, leaving the algorithm to search for a corrected solution to the model. In MoMA, the given feedback is incorporated into the ILP via additional constraints. Using warm-starts allows optimizing the modified problem fast enough for interactive use. Fixing a single error will usually resolve a bulk of transitive errors. These interaction-based modifications and re-optimizations are iterated until the found solution is satisfactory to the user, i.e., appears to be free from errors.

Below we will introduce the five specific interaction primitives that MoMA allows for. We will see that all these editing primitives do not require significant changes to the existing tracking software and can be implemented very efficiently. To illustrate how interactive curation works in practice, we provide an [introductory movie](https://github.com/fjug/MoMA/wiki/movies/MoMA_LeveragedEditing.mp4) that shows several of these primitives in action.

#### 2.2.1 Forcing Segments

One possible error is that the tracking may have failed to identify a particular cell, possibly across multiple frames. In this case, the user can hover the mouse over the part of the image where a cell was not picked up by the original optimization, and inactive segment hypotheses underlying that part of the image will be highlighted. Clicking on any highlighted segment will cause the system to (*i*) modify the ILP as described below, and (*ii*) re-run the solver to obtain the optimal solution for the given data, now constrained to include the forced segment.

Technically this can be achieved by adding a single constraint to the ILP, namely *h* = 1 where *h* is the chosen segment. Applying the redundancy reduction discussed at the end of Section 2.1, the constraint to be added can be expressed in terms of assignment variables as

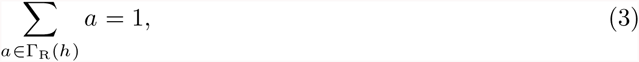

where Γ_R_(*h*) is the right neighborhood of *h*, i.e. the set of all assignments leaving *h* towards the next frame.

#### 2.2.2 Excluding Segments

In addition to allowing users to force missing segments to be included, the user can also force MoMA to exlude certain segments from its solution. The re-solved ILP will correspond to the minimal cost solution for the data, constraint to exclude the chosen segment. Analogously to forcing segments, the constraint to be added to the ILP is

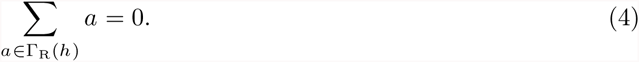

#### 2.2.3 Forcing Assignment

Instead of interacting with segments, a user might want to directly work with individual assignments. To do this, the user can browse through a library of available assignments. Any such assignment can be chosen to be included in future tracking solutions. Note that this also enforces the solution to contain specific segments, namely the segments that are associated with the forced assignment.

Browsing the library of available assignments can be done in only a few mouse-clicks. Since there is precisely one binary variable *a* corresponding to the chosen assignment, the respective constraint to be added to the ILP to force this assignment to be 1 (on) is simply *a* = 1.

#### 2.2.4 Excluding Assignments

Analogously, the user might want to exclude an assignment that is part of the current tracking solution. A single click suffices to exclude a particular assignment. The corresponding constraint to be added to the ILP for excluding an assignment is simply *a* = 0, where *a* is the chosen assignment.

#### 2.2.5 Specifying the Number of Visible Cells

For all previous interactions the user had to pick individual assignments or segments to include or exclude. The last interaction primitive that MoMA includes operates on a semantically higher level.

One or multiple errors can often be fixed by telling the model how many cells are contained in the current time-point. Single missing or superfluous segments can be added or removed, but erroneous or missing divisions also be undone or forced. In extreme cases, with multiple errors occurring at a single time point, one hint by the user can suffice to resolve all problems at once. As before, let us constrain the solution space in the way we want – here to only those solutions that contain *k* segmented cells at time-point *t*. Formally this is accomplished by adding the constraint

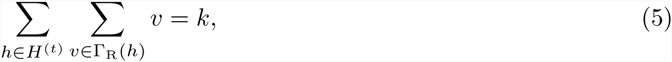

where *H*^(*t*)^ is the set of all segments existing at time *t*.

### 2.3 Implementation, Data Export, and Extension Modules

MoMA is implemented in Java, using the imaging library ImgLib2 [32] and other components from the open source universe around ImageJ and Fiji [33, 34]. For solving ILPs we use Gurobi™. The source code of MoMA is a Maven project, hosted on GitHub (http://github.com/fjug/MoMA). The same page also hosts a comprehensive Wiki containing information about how to install and use MoMA.

Addutional useful features of MoMA include (*i*) the ability to optimize (solve) only parts of a loaded dataset, (*ii*) save a fully or partially curated dataset, and (*iii*) the possibility to export a found tracking solution for downstream processing. Below we will discuss these issues in detail.

#### 2.3.1 Interactive Editing of Large Datasets

If a loaded dataset contains 1000 or even more time-points, the re-optimization of MoMA’s assignment model can take tens of seconds. While this is still very fast, e.g.when compared to the data acquisition time for such a dataset, leveraged editing can become cumbersome when the user is forced to wait tens of seconds between interactions for the re-optimization to finish. In order to guarantee fast interactive response times, MoMA allows users to define a subrange of time-points {*t*_*a*_, *t*_*b*_} for which to perform the optimization.

All assignments that are not in {*t*_*a*_, *t*_*b*_} are either set to the value computed at a previous (partial) optimization run, or simply clamped to be 0. Formally this can be expressed by

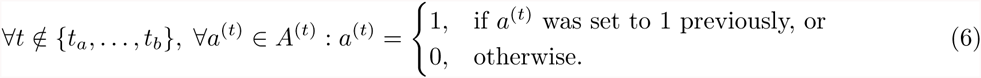

#### 2.3.2 Saving and Loading Curated Datasets

In practice it is important that users can save and re-load the curations that they performed. As we have seen in Section 2.2, each editing primitive corresponds to introducing an additional constraint to the underlying optimization problem. MoMA is capable of serializing all executed user edits in a simple, human readable file. The saved state can easily be loaded by first opening the same raw dataset and then loading such a previously serialized file.

#### 2.3.3 Data Export and Data Formats

MoMA data export can be modified based on the data a user wants to export. An exhaustive list of exportable data is given below. Users should check the MoMA Wiki for detailed information regarding the data format (https://github.com/fjug/MoMA/wiki).

- Data source.
- Total number of cells observed in the dataset.
- Number of channels in the raw data, i.e. phase contrast and fluorescent channels.
- Growth channel (GC) height and image height in pixels.
- (Vertical) position of the GC in the image.
- For each cell, its cell id, and lineage information (the ids of its ancestors)
- For each time point in the life of each cell: position in the GC [pixels and cell number]; cell length; bounding box area; intensity histogram, intensity percentiles, and pixel intensities for the all channels.

## 3 Using MoMA and post-processing its data

### 3.1 Loading time and export

All the curating with MoMA was performed on a MacBookPro (2.4 GHz Intel Core i7, 8GB of memory). On this setup loading, initialisation and the first round of optimization of a dataset with 480 frames with two channels typically takes around 1 minute. After curating the data the export step takes another 30 seconds.

For the analysis of the glucose/lactose experiments for which we show data in this report we selected GCs that are well filled with cells in the beginning and still harbor cells at the end of the experiment. GCs in which most cells get lost and only few cells move around in the GC were also excluded. We made sure only to include GCs that showed no structural defects and furthermore we excluded GCs that harbored cells that lysed or showed abnormal elongation.

### 3.2 Curation times and performance

MoMA was tested on Mother Machine data with ~30 frames per cell cycle, stable focus over the experiment and two different channels imaged. To estimate the time the user needs to spend to curate the datasets, an unbiased selection of GCs was analyzed and the time spend for curating was measured. For the selection of the GCs there was no visual inspection of the GCs other than checking that they harbor cells on the first frame. Therefore this sample also harbored GCs in which the cells are lost during the experiment or some that show structural defects. For the curation times shown in the paper we only used the times for the GCs in which we had cells till the end of the experiment to make sure curation times are not underestimated by including GCs in which only a small fraction of the frames actually shows cells. Defect GCs were excluded as well. There are also some GCs in which a cell is lysing or shows other abnormalites. In such cases even with the eyes of an experienced observer it is difficult to decide on the border of such cells and here also MoMA struggles since such cells show completely different intensity profiles and behave differently so that the cost functions are not applicable anymore.

### 3.3 Refinement of the cell size estimation

From the imaging data we obtain, for each cell, pictures for each time point during its life-cycle. The software does not determine the precise cell outline but instead a rectangular bounding-box within wich the cell is fully contained. An example of such a phase contrast picture is shown in Fig. 6. The cell is shown in darker colors.

**Figure 6:**
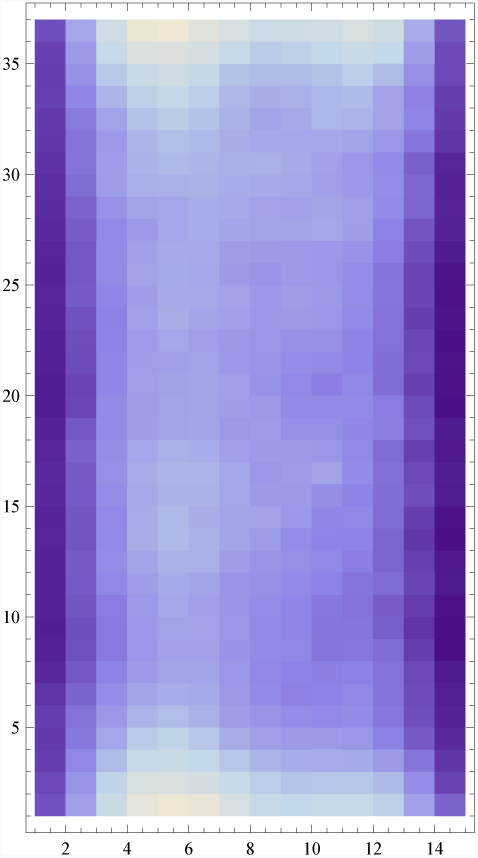
Example phase contrast picture of a rectangular image segment containing a single cell. The cell corresponds to the region of lower intensities.

In our cell size estimation we estimate the size of the cell by estimating which *pixel rows* correspond to the top and bottom of the cell. To this end we sum the pixel intensities in each row of the phase contrast image and analyze the profile of row-sums. Roughly speaking, the rows with low row-sums correspond to the cell, and the edges at the top and bottom of the box that have high row-sums are outside of the cell. Let *r*_*i*_ denote the sum of intensities of row *i*. The general shape of the profiles *r*_*i*_ as a function of *i* is that the row-sum is low in the middle of the cell and increases at the ends of the cell. Four examples of row-sum profile (the same cell at 4 different time points) are shown in the left panel of Fig. 7.

**Figure 7:**
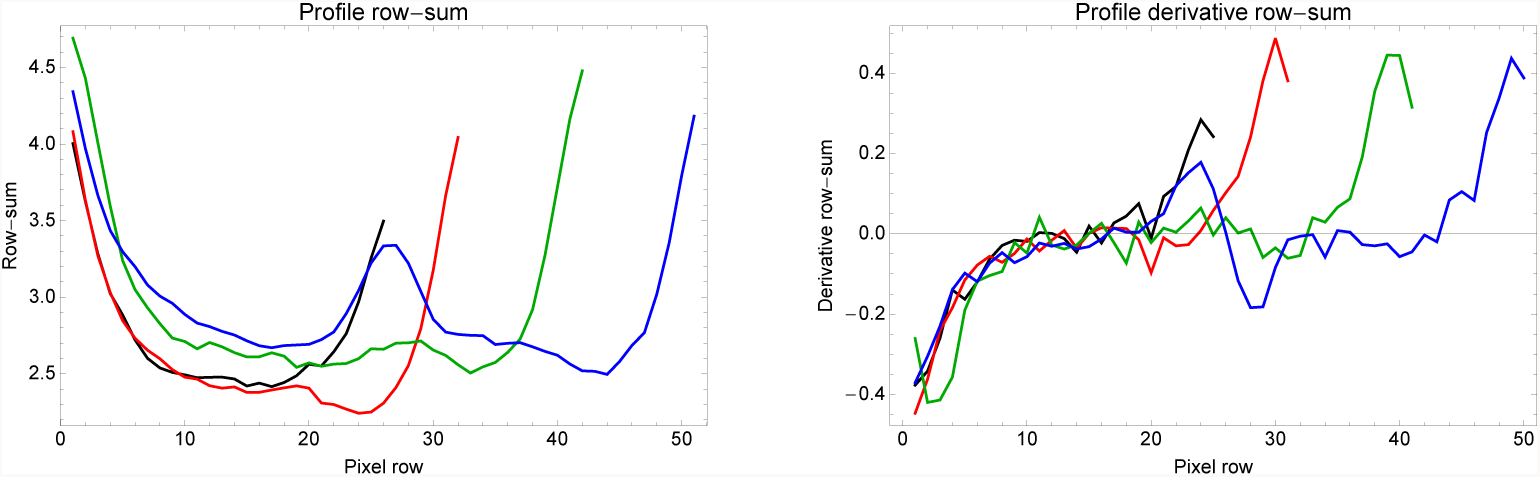
Left panel: Four examples of the row-sum profiles *r*_*i*_, i.e. sum of the phase contrast pixel intensities along a pixel row as the rows go from the bottom to the top of the rectangle in which the cell is contained. The four colors correspond to the same cell at time points 1 (black), 7 (red), and 14 (green), and 22 (blue), which is the last time point before division of this cell.**Right panel**: Profiles of the derivatives of the row-sums for the same 4 time points.

The absolute heights of the profile vary substantially along the cell cycle so that it is not straightforward to determine the cell’s edges from the row-sum profile directly. However, we noticed that the *derivatives* of the row-sums, i.e. *d*_*i*_ = *r*_(*i*+1)_ − *r*_*i*_, are more consistent for the same cell at different time points (right panel of Fig. 7). However, even these profiles cannot be counted on to have a uniform shape across cells and time points. Notice, for example, that just before division of the cell, there is a local maximum of the row-sums in the middle of the cell.

After experimenting with a number of different criteria, we settled on the following simple procedure:

1. Define the **middle of the cell** *i*_mid_ by finding the ten consecutive rows with lowest total row-sum, i.e. such that the sum 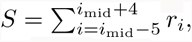 is minimal. The number 10 was picked to be large enough to effectively average out fluctuations, while still being small enough to ensure its short relative to the total length of the cell.
2. The **left end** *i*_*l*_ of the cell is defined as the row to the left of the middle, which has *smallest* derivative, i.e. *d_il_ ≤ d*_*i*_; *∀i < i*_mid_.
3. The **right end** *i*_*r*_ of the cell is defined as the row *i* that lies to the right of the middle and has the *largest* derivative, i.e *d_ir_ ≥ d_i_ ∀i > i*_mid_.
4. The size of the cell is defined as the difference between the left and right-end: *s* = *i_r_ − i*_*l*_.

As we show below, with this procedure virtually all cells accurately follow simple exponential growth curves as a function of time, supporting the robustness of the procedure. However, we found that for bottom cells this procedure breaks down when the bottom cell is pushed against the bottom of the GC. For such cells there is no maximum in the derivative *d*_*i*_ at the right end of the profile, but *d*_*i*_ instead drops at the right end of the profile, which corresponds to the bottom of the GC. For these cells we estimate their bottom end by finding the smallest *d*_*i*_ to the right of the middle. This alternative procedure introduces a systematic over-estimation of the length of bottom cells whenever they touch the bottom of the GC. To correct for this we subtract a constant number of pixels *db* from touching bottom cells, and we set *db* such that, in constant environmental conditions, the fit of the growth-curves to exponentials is maximized. Finally, our procedure for finding the top and bottom of the cell may introduce a systematic offset to the true size of the cell. To correct for this we added an additional size *ds* to each cell’s estimated length and fitted *ds* so as to maximize the overall fit of the growth-curves to exponentials for the corrected cell sizes.

#### 3.3.1 Performance of the size estimation

It is clear that our cell size estimation is quite coarse and we wanted to get some idea of how accurate our estimates are. This is difficult to do directly because we do not have independent measurements of cell sizes that can be used as a gold standard. However, if we find that the estimated cell size *s*(*t*) accurately follows a simple exponential or linear form as a function of time *t*, then this suggests the errors in cell size are at most as large as the fluctuations of *s*(*t*) away from the simple exponential or linear growth law.

Let *s*(*t*) be the estimated size of the cell at time *t* and *x*(*t*) = log[*s*(*t*)]. For each cell, we calculate the Pearson correlation between *x*(*t*) and *t* across the cell cycle, as well as the Pearson correlation between *s*(*t*) and *t*. Figure 8 shows the cumulative distributions of the Pearson correlation of the growth curves with exponential (red) and linear (blue) functions for cells grown in lactose (left panel) and in glucose (right panel).

**Figure 8:**
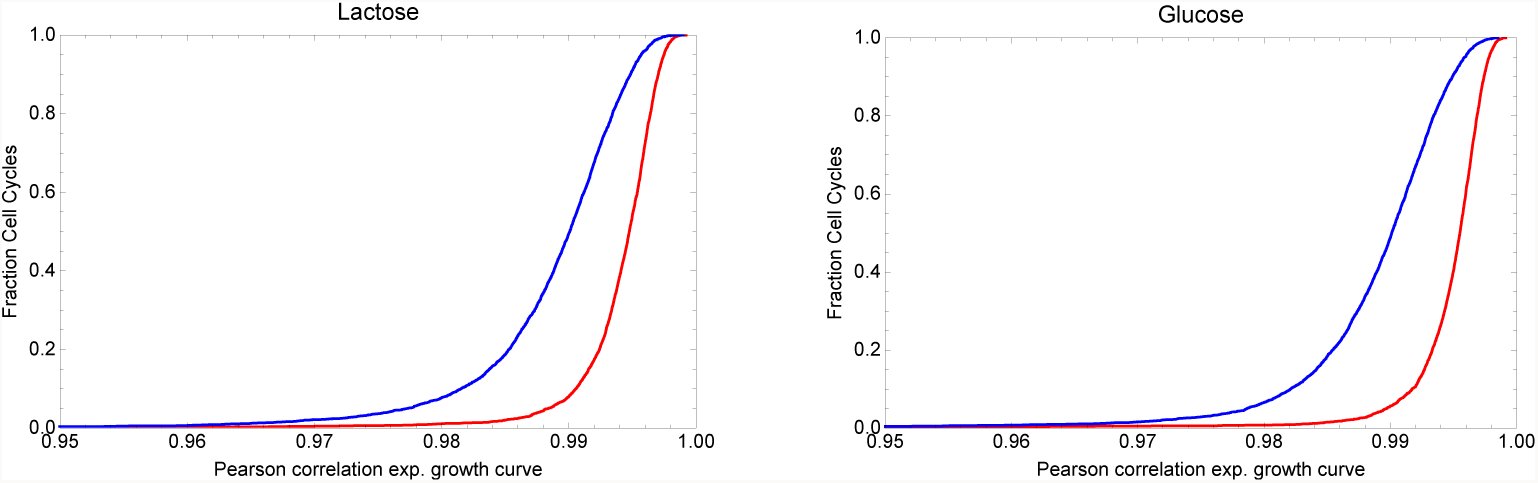
Cumulative distributions of the Pearson correlations between the estimated sizes of the cells and time (blue), or estimated log-sizes and time (red) for the cells grown in lactose (left panel) or glucose (right panel).

We see that the growth curves are very well described by exponential functions of time, i.e. the median correlation coefficient is approximately 0.995 and almost all cells have correlation coefficients larger than 0.99. Correlation coefficients are substantially lower for fits to linear growth curves. Note that, whereas correlation coefficients are still very high for the linear growth fits, the log-likelihood for a growth-curve with correlation *r* and *T* time points scales as −*T* log[1 − *r*^2^]. Thus, for a growth-curve with *T* = 30 time points, the log-likelihood difference *dL* between a fit with *r* = 0.99 and one with *r* = 0.995 and 0.99 is *dL* = 20.7.

### 3.4 Estimation of fluorescence per cell

To estimate the GFP content of each cell we post-process the fluorescence data as follows. The raw data consists of fluorescence intensities for all pixels within the segment of the picture containing the cell. This segment is 100 pixels wide, with the GC covering approximately 13 pixels in the center of the picture. We first obtain *column-sums c*_*i*_ by summing the pixel intensities of all pixels in each of the 100 columns *i*. Note that we assume that these column sums are dominated by the fluorescence coming from the cell in question, i.e. that the fluorescence coming from neighboring cells above and below the cell are negligible. We find that this is a good approximation when cells in a given GC all have similar fluorescences but note that, in conditions where neighboring cells may have fluorescences that differ by orders of magnitude, this assumption may break down. Figure 9 below shows the profile of column sums *c*_*i*_ for a cell at three time points of its cell cycle while growing in lactose (top three panels) and for a cell growing in glucose (bottom three channels). From prior biological knowledge, we know that GFP is highly expressed during the growth on lactose, and that it is essentially not expressed (or very lowly expressed) during the growth on glucose.

**Figure 9:**
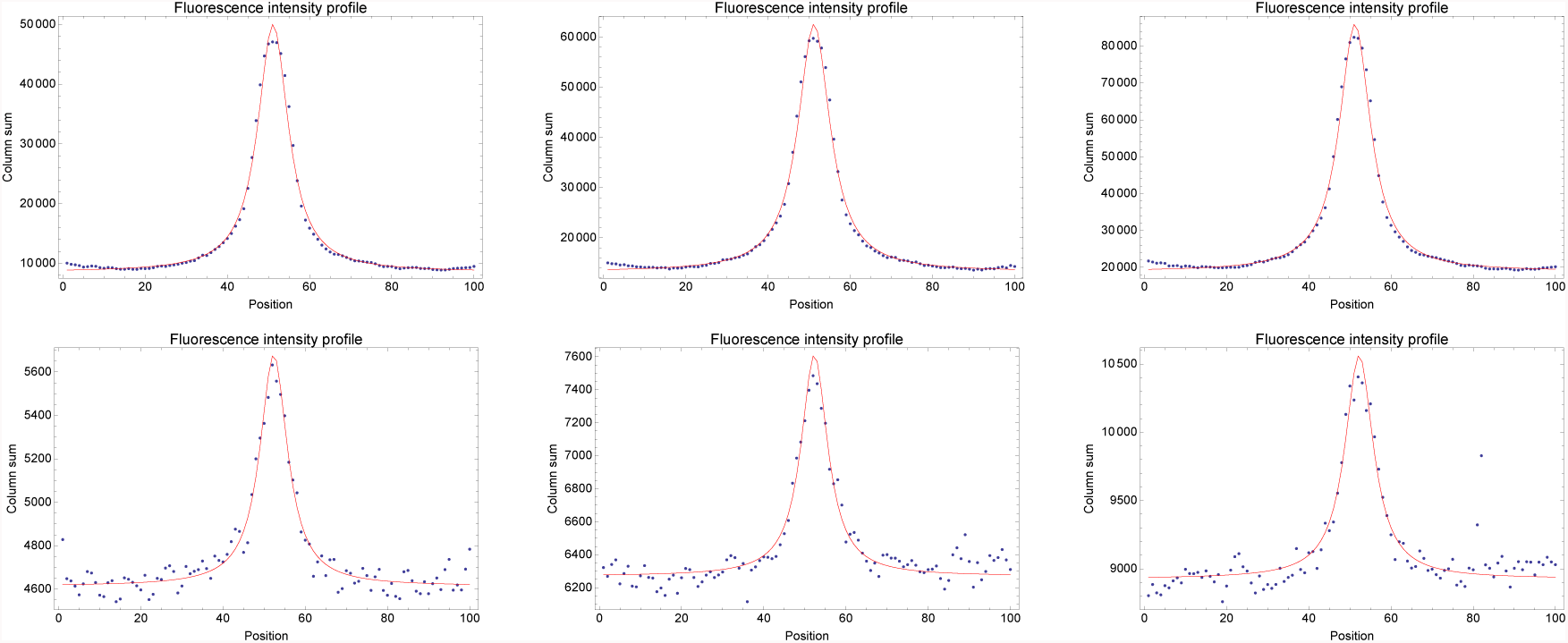
Examples of the horizontal profiles of column sums *c*_*i*_ of fluorescence intensities (blue dots) for three cells during growth on lactose (top panels) and during growth on glucose (bottom panels). The three panels show, from left to right, the profiles at the start, the middle, and at the end of the cell cycle of two example cells. The red curves show the fits to the mixture model of a fixed background plus a Cauchy distributed signal.

Remarkably, in contrast to phase contrast images, the GC (central 13 pixel positions in the figures) is not detectable at all in the fluorescence curves. Instead, the fluorescence signal shows a long-tailed peak centered in the middle of the GC, extending far beyond the width of the GC, and reaching a minimum at positions halfway between neighboring GCs, i.e. near the left and right ends of the profiles in Fig. 9. As the cell grows, i.e from the leftmost to rightmost panel, the length of the segment grows and the column-sums grow proportionally to the segment length. Notably, this minimal fluorescence level is almost twice as high when growing in lactose compared to when growing in glucose. We conclude from these observations that the fluorescence from each cells spreads over significant distances across the image and that this also causes background levels to depend on the overall expression levels in neighboring GCs. Therefore, to properly estimate the amount of fluorescence emerging from the cell we need to fit the background intensity within each segment and we need a mathematical model for the long-tailed shape of the peak.

We found that the shape of the peak is very well described by a Cauchy (or Lorentzian) distribution, giving an overall form of the fluorescence profile:

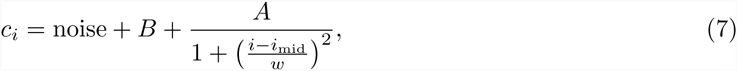

where *i* is the horizontal position, *i*_mid_ is the center of the signal, *w* its width, *B* the background fluorescence, is *A* the amplitude of the signal, and ‘noise’ is the measurement noise. Assuming that the measurement noise is Gaussian distributed, it is straight forward to fit this model using expectation maximization. We find that, systematically, the center *i*_mid_ *≈* 50 − 52, and the width *w* ≈ 5 − 6 pixels. We will interpret the amplitude *A* to be proportional to the total number of GFP molecules in the cell. The expectation maximization procedure for fitting the fluorescence profile is

1. Find the maximum and minimal fluorescence column-sums *c*_max_ and *c*_min_ across the profile.
2. Initialize *B* to *c*_min_, *A* to *c*_max_ − *c*_min_, *w* to 5.5 and *i*_mid_ to 50, i.e. in the middle of the profile.
3. Calculate a theoretical profile:

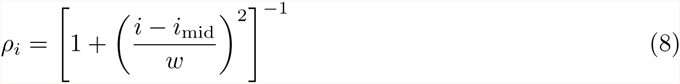

and its integral 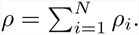
4. Set a new value of the background *B*′:

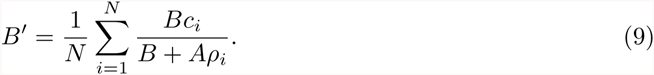
5. Set a new value for the amplitude *A*′:

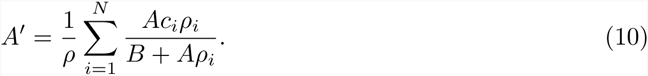
6. Using the updated values *A*′ and *B*′, calculate an updated profile *ρ*_*i*_ and optimize *i*_mid_ by finding the zero of the derivative:

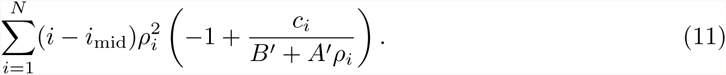
7. Using the updated values *A*′, *B*′, and *i*_mid_, optimize *w* by finding the zero of the derivative:

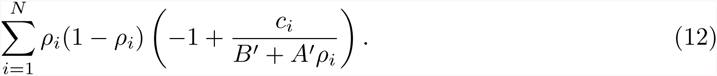

### 3.5 Data analysis methods

#### 3.5.1 Estimating the cell’s autofluorescence

To estimate the autofluorescence of the cells, i.e. in addition to the background fluorescence of the medium and PDMS, we measured the wild-type strain of E. coli MG1655, i.e. without the fluorescent reporter, in the conditions where we switch between glucose and lactose. We observed that the estimated total fluorescence, i.e the amplitude *A* from the previous section, correlates well with the size of the cells during their cell cycle. That is, i.e. fitting a linear relationship *A* = *aS* + *b* of the fluorescence *A* as a function of the estimated cell size *S* typically yields Pearson correlation coefficients of *r* ≈ 0.9. Moreover, we observed that the vast majority of fits were consistent with *b* = 0, i.e. the total fluorescence being directly proportional to cell size, supporting that this signal corresponds to the autofluorescence of the cell.

To fit the autofluorescence *a* (per micron of cell length) we selected all cells that were observed for a full cell cycle, who never got within 100 pixels of the end of the GC during their cell cycle, and whose length as a function of time was well fit by a simple exponential growth curve ((*r*^2^ *≥* 0.99). This latter restriction mainly serves to remove cells that had a transient stop in growth after the first switch to lactose. In total there were 284 cells that passed all these criteria. For each of these cells we replaced the directly estimated sizes *S*_*t*_ at each time point *t*, with the sizes 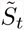 estimated from the exponential fit of *S*(*t*) as a function of time (reasoning that these estimates are more accurate than the direct measurements). For each cell we then fit a function 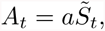 assuming Gaussian measurement noise of unknown variance.

That is, for a single cell we write

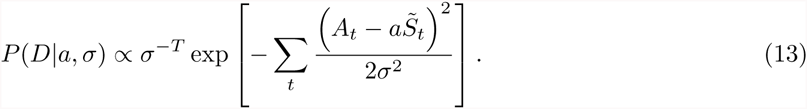

Using a scale prior on *σ* of the form *P* (*σ*) *∝* 1*/σ*, and integrating over *σ* we obtain

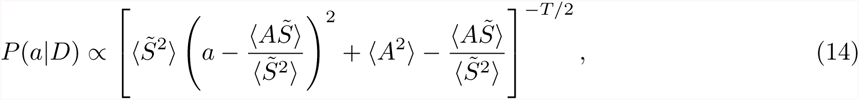

where *T* is the number of time points in the cell cycle and the averages are over the time points in the cell cycle.

The optimal value of *a* is given by

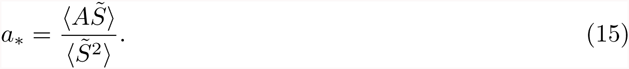

Approximating the posterior by a Gaussian we obtain for the standard-deviation of the estimated *a*

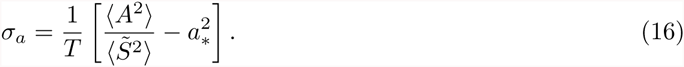

Figure 10 shows the estimated value *a*_*∗*_ and its error bar *σ*_*a*_ for each of the 284 cells (left panel). Note that, although most cells have fluorescence values between 400 and 500, there are some outliers at higher fluorescence. This is also evident from the combined probability density of *a* values (right panel).

**Figure 10:**
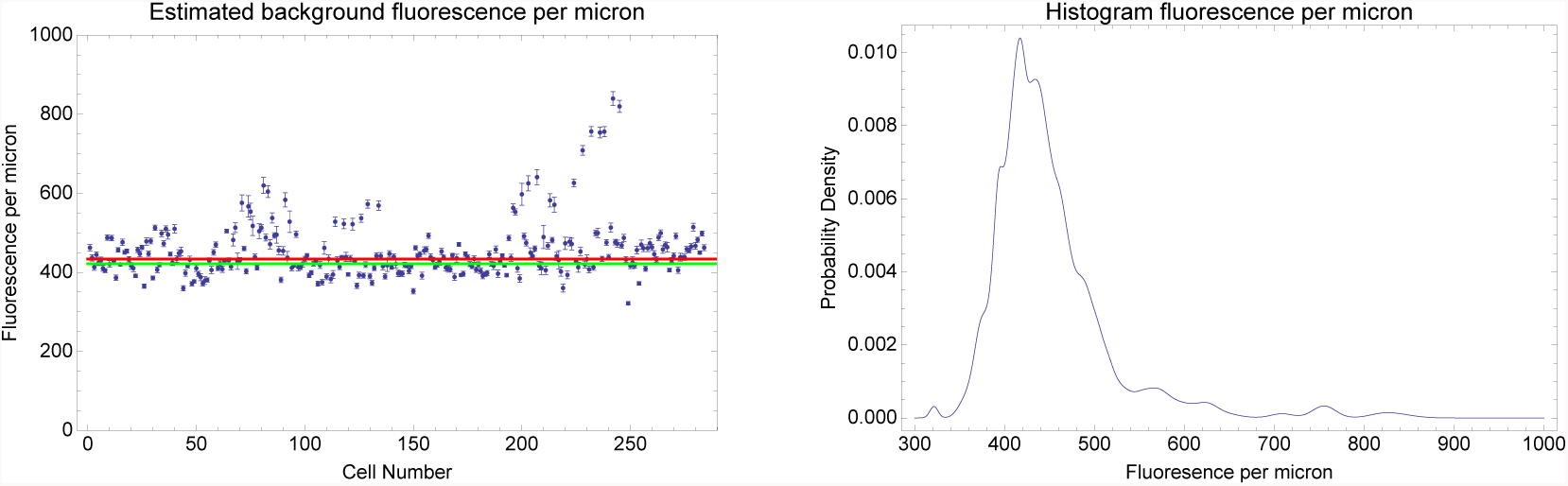
**Left panel**: Estimated background fluorescence per micron cell length *a*_*∗*_ and its error bar *σ*_*a*_ for each of the 284 cells for which fluorescence was fit as a function of cell size. The red line (*a* = 433.5) is the fit obtained when all cells are assumed to have a common fluorescence per micron *a*. The thick green line is obtained with a mixture model that allows for ‘outliers’ from a uniform distribution (*a* = 421.8). **Right panel**: The joint probability density of *a* given by the mixture of Gaussian distributions for all 284 cells.

If we assume there is one common background fluorescence per micron *α* for all cells, then the probability of the data given *α* is given by

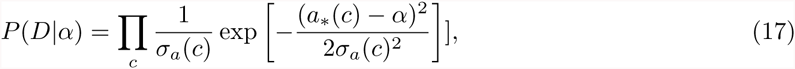

where the product is over the 284 cells *c*.

Maximizing this function with respect to *α* yields

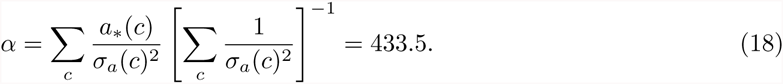

If we allow that there are some ‘outlier’ cells whose value of *a* is described by a uniform distribution of width *W* = *a*_max_ − *a*_min_, then the likelihood of the data as a function of *α* and the fraction of non-outlier measurements *ρ* is given by

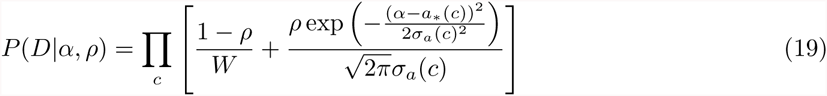

Maximizing this function with respect to *α* and *ρ* yields *α* = 421.8 and *ρ* = 0.31. In the following we will use this latter value of *α* for the autofluorescence per micron of cell length. For each cell with estimated size *S* and total fluorescence *A*, we thus obtain an autofluorescence corrected fluorescence level 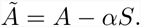.

#### 3.5.2 Estimating the rate of bleaching and degradation of GFP

As shown in Figure 2A, while the lac operon is induced in the lactose phases, GFP production ceases during the glucose phases. In this regime, the total cell fluorescence slowly decreases during the cell cycle, and approximately divides in half at cell division. We reasoned that the decay of fluorescence during the cell cycle is the result of GFP bleaching and, potentially, also some GFP degradation. Inspection of the data indeed shows that the total fluorescence decrease is captured better by an exponential rather than linear model. For this analysis, we consider only observations between 30 min after the switch to glucose and before the next switch to lactose, and to cells with at least 10 points in this time window. Although the Pearson correlation of the fits doesn’t favor one model over the other (which can be explained by the narrow variations of GFP levels within each cell cycle, typically less than 20%), comparing the fitted slopes is instructive: while linear slopes strongly depend on the absolute GFP level of the cell, exponential rates are constant over all the range of GFP levels (Fig. 11).

Hence assuming that a cell undergoes bleaching + GFP degradation at a rate *µ* per second, we fitted *µ* in each cell as minus the slope of a linear model of time and log(GFP level). Using individual rates and their standard deviations, we estimate the global rate *µ*_*∗*_ to be equal to 5.3 × 10^−*5*^ ± 5.10^−7^ (mean *±* sd).

#### 3.5.3 Inferring the conversion factor between fluorescence and GFP number

To estimate the conversion factor between total fluorescence 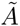 and the number of GFP molecules we will use data on the fluctuations in fluorescence levels of newborn sibling pairs. To avoid confounding effects from GFP production, we collected division events from the glucose phases in our switching experiments, when GFP production has ceased. Collecting division events from these phases has the added advantage that absolute GFP levels vary over a considerably range across cells during these phases, allowing us to quantify the size of fluctuations in sibling fluorescence as a function of total fluorescence. Our observations consist of fluorescence levels at birth (*x_i_, y*_*i*_) for sibling pairs of daughters, where *i* runs from 1 to *N*, with *N* the total number of such sibling pairs. Using the same criteria as in the decay analysis for mothers and daughters, we collected *N* = 357 sibling pairs. GFP levels at birth were estimated in each daughter cell as average of the levels at all time points corrected for the previously estimated decay. Assuming that the GFP molecules in the mother cell are distributed randomly between the daughters, the fluctuations in the numbers of GFP molecules going to each daughter should be binomial distributed, and this has been used previously to infer a conversion factor between GFP molecule numbers and fluorescence levels [18]. In particular, assuming binomial fluctuations, the expectation of the square of the difference 〈(*n_i_ − m*_*i*_)^2^〉 should be equal to the total count *n*_*i*_ + *m*_*i*_. Given an conversion factor *λ*, such that the GFP molecule counts correspond to (*n_i_, m*_*i*_) = *λ*(*x_i_, y*_*i*_), one can estimate *λ* by observing

**Figure 11:**
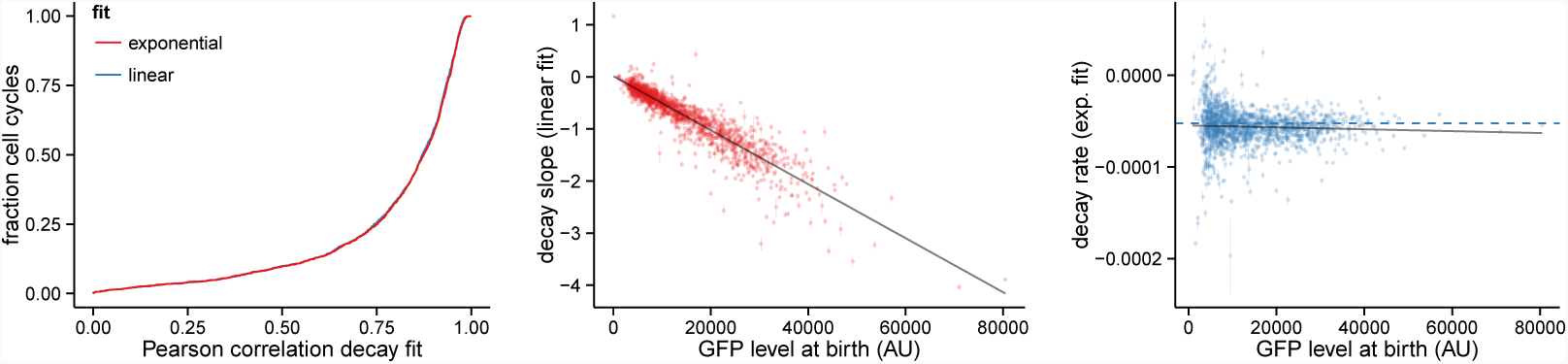
Analysis of the GFP decay based on cells observed after the lactose-glucose switches when no GFP is synthesized. **Left panel:** Cumulative distributions of the Pearson correlations between the estimated GFP levels and time (red), or estimated log-GFP levels and time (blue). **Center panel:** using a linear model to describe the decay, the slopes in individual cells depend strongly on the total GFP levels in the cell. **Right panel:** using an exponential model, the rates are independent of the total GFP levels in the cell (dashed line shows the average rate used in subsequent analysis).

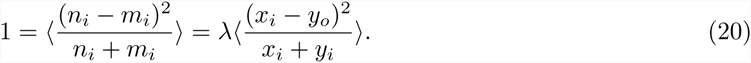

However, using this ‘naive’ approach, we find that the conversion factor *λ* systematically decreases with total fluorescence (Fig. 12, left panel), changing by as much as fourfold depending on whether division events with low or high absolute fluorescence are used. This result implies that the variance of fluorescence fluctuations grows faster than linear with total fluorescence, suggesting that there are additional fluctuations with variance proportional to total fluorescence squared. Inspection of the data strongly suggests that these additional fluctuations derive from fluctuations in the *cell size* of the daughters. That is, in addition to the binomial fluctuations there are fluctuations caused by the daughters having unequal size. Practically, cell size at birth is estimated in each daughter cell by extrapolating the linear fit of time and log(length). Indeed, we observe a substantial correlation between the relative sizes of the siblings and the relative amounts of fluorescence each sibling receives (Fig. 12, right panel, Pearson correlation *r* = 0.44)

We thus developed a more sophisticated model, which takes into account fluctuations in the cell sizes, the binomial fluctuations, as well as measurement noise. For a given division event *i*, let *ρ*_*i*_ denote the measured fraction of the cytoplasm that went to the first daughter, and let *q*_*i*_ = *x*_*i*_/(*x*_*i*_ + *y*_*i*_) be the measured fraction of the fluorescence that went to the first daughter. We will assume that *q*_*i*_ is a noisy measurement of the true fraction of molecules *q* = *n*_*i*_/(*n*_*i*_ + *m*_*i*_) that went to the first daughter, and that *ρ*_*i*_ is a noisy measurement of the true fraction of the mother’s cytoplasm *ρ* that went to the first daughter. Given *ρ* and a total number of molecules *n* = (*n*_*i*_ + *m_i_*), the molecule numbers (*n_i_, m_i_*) will show binomial fluctuations and the fraction *q* will have a variance var(*q*) = *ρ*(1 − *ρ*)*/n*. In addition to this variance we will assume there is a total measurement noise of variance *v*, so that the total expected square-deviation between the measurements *q*_*i*_ and *ρ*_*i*_ should be *v* + *ρ*(1 − *ρ*)*/n*. We will assume that the sum of these fluctuations due to the binomial noise and measurement noise is approximately Gaussian distributed. Finally, we will assume that the binomial variance *ρ*(1 − *ρ*)*/n* is well approximated by the measured values *ρ*_*i*_(1 − *ρ*_*i*_)/(*λ*(*x*_*i*_ + *y*_*i*_)).

**Figure 12:**
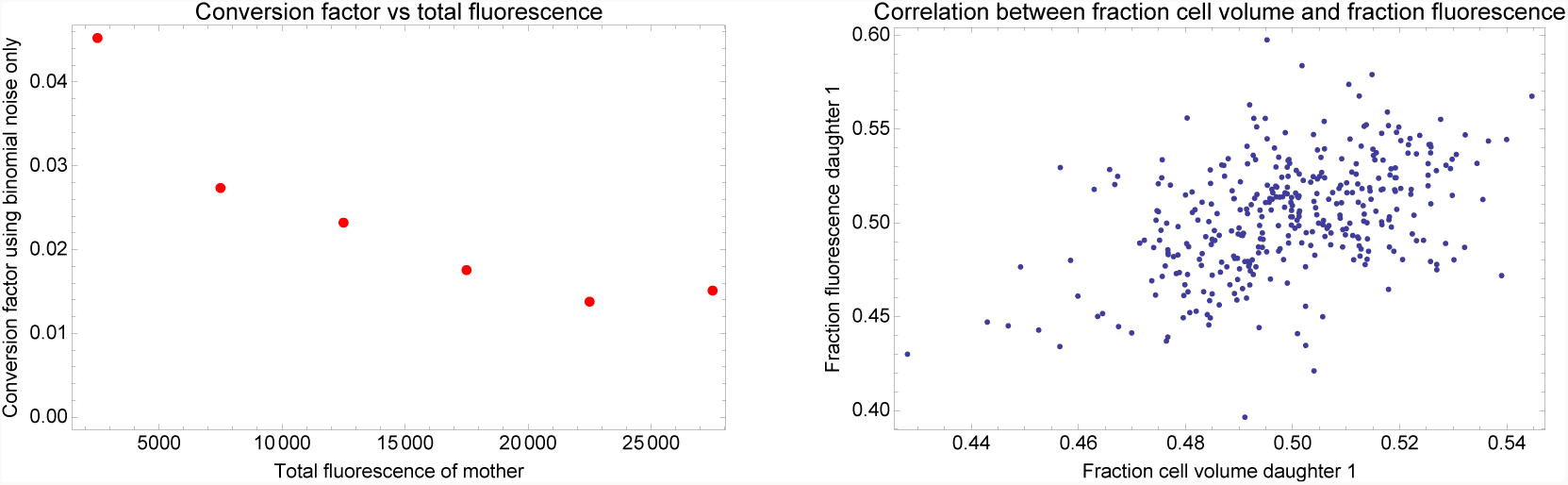
**Left panel**: Estimated conversion factor *λ*, using the ‘naive’ method which assumes there are only binomial fluctuations, as a function of total fluorescence. Division events were divided into bins based on the mother’s total fluorescence. Binsize was 5000. For each bin, the conversion factor *λ* was estimated assuming only binomial noise. **Right panel**: Correlation between fluctuations in fluorescence and cytoplasm size of two sibling cells immediately after birth. Each dot corresponds to a pair of sister cells with the horizontal axes showing the fraction of the total cytoplasm that is taken up by the first sister, and the vertical axis showing the fraction of the total fluorescence taken up by the first sister.

Under this model, the probability of observing the fraction *q*_*i*_, given the measured volume fraction *ρ*_*i*_, the conversion factor *λ*, and the total measurement noise *v* is given by

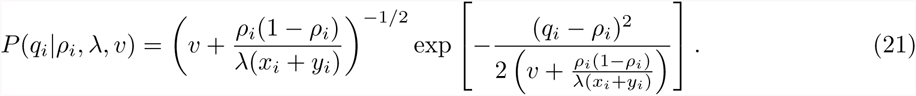

Summing over all *N* division events, the log-likelihood of *λ* and *v* is now given by

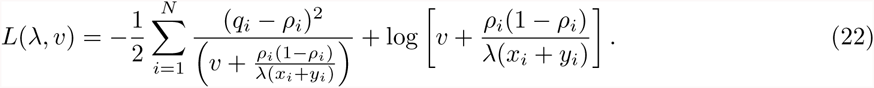

To obtain the posterior probability of *λ* we marginalize over the unknown variance *v* (using a uniform prior). That is, we calculate *L*(*λ*) = log ^Ir^ exp[*L*(*λ, v*)]*dv*^r^. Using this model, the maximal likelihood value of *λ* is given by

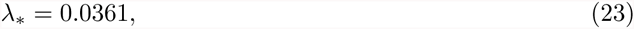

and the symmetric 95% posterior probability interval is given by *λ* ∈ [0.026, 0.112].

Figure 13 shows the posterior distribution *P* (*λ|D*) obtained with our model. For comparison, Fig. 13 also show the conversion factors that would be obtained with the naive method that assumes there is only binomial noise.

#### 3.5.4 Estimating lag times to *lac* operon induction and shut down

In Figure 2 we show results from experiments where the carbon source is alternated between glucose and lactose every 4h with the *lacZ-GFP* strain. As expected the expression of *lacZ-GFP* is induced in each phase of the experiment with lactose present. To estimate the time until the *lac* operon is induced after a glucose-lactose switch in a given cell, we consider only cells that are born before the switch and last for at least 3 frames. We checked visually that no induction happens in the first 10 min in all three switches, hence the pre-switch GFP level is estimated as the average of the first 3 time points. Since GFP levels are increasing monotonically at the start of each lactose phase, we measured the delay until the GFP level has increased by a fixed amount, here 200 molecules. This delay is interpreted as the induction time of the *lac* operon plus the maturation time of the *lacZ-GFP* fusion proteins.

**Figure 13:**
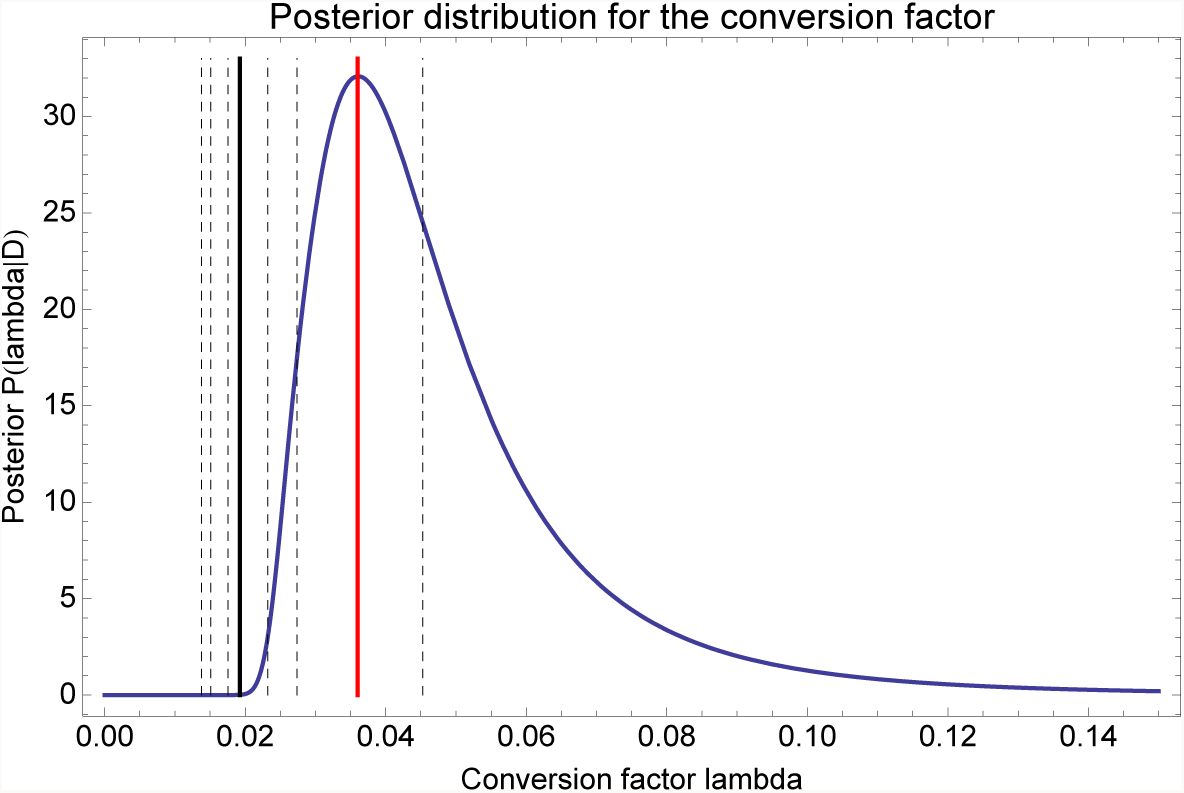
Posterior distribution *P* (*λ|D*) for the conversion factor *λ* given the data on our sibling pairs (blue curve). The red line shows the maximum likelihood valeu *λ*_*∗*_ = 0.0361. The black lines show the estimated conversion factors that are obtained when assuming binomial noise only. The solid line results from using all data, and the dashed lines result from different subsets at different absolute fluorescence values (see Fig. 12).

Similarly, to estimate the time until the GFP signal decreases at the lactose-glucose switches, we selected cells that are born before the switch and persist in the channel for at least 1h after the switch. For each cell we measured the delay between the switch and the time when the GFP level reaches its maximum. We use the measured time as an estimation of the time needed to switch off the expression from the *lac* operon plus the maturation time of the *lacZ-GFP* fusion proteins.

